# Language impairment with a partial duplication of *DOCK8*

**DOI:** 10.1101/2020.06.16.155523

**Authors:** Antonio Benítez-Burraco, Maite Fernández-Urquiza, Mª Salud Jiménez-Romero

## Abstract

Copy-number variations of the distal region of the short arm of chromosome 9 are associated with learning disabilities and behavioral disturbances. Deletions of the 9p are more frequent than duplications. We report in detail on the cognitive and language features of a child with a duplication in the 9p24.3 region (arr[hg19] 9p24.3(266,045-459,076)x3). He exhibits marked expressive and receptive problems, which affect to both structural aspects of language (notably, inflectional morphology, complex syntax, and sentence semantics), and to functional aspects (pragmatics). These problems might result from a severe underlying deficit in working memory. Regarding the molecular causes of the observed symptoms, they might result from the altered expression of selected genes involved in procedural learning, particularly, some of components of the SLIT/ROBO/FOXP2 network, strongly related to the development and evolution of language. Dysregulation of specific components of this network can result in turn from an altered interaction between DOCK8, affected by the microduplication in 9p24.3 borne by our proband, and CDC42, acting as the hub component of the network encompassing language-related genes. Still, some genes found strongly upregulated in the subject and not related to these genes, particularly *NRCAM*, can contribute to the observed problems in the language domain, as well as to specific features of the proband, particularly, his impulsivity.

## INTRODUCTION

Copy number variants (CNVs) identified in subjects with language problems provide important evidence of how changes in gene dosage affect the wiring and functioning of brain areas involved in language processing, particularly, if they result in syndromic features and encompass few genes. The terminal region of the short arm of chromosome 9 is prone to deletion, giving rise to a complex syndromic condition entailing mental retardation, delayed psychomotor development, and speech delay (OMIM#158170). Specifically, and according to the Unique database (https://www.rarechromo.org/), deletions of the 9p24.3 region are associated with mild learning disabilities, that commonly improve with appropriate early/supporting intervention, and with language and speech delay, with a greater impact on the expressive domain. Although deletions of the 9p are more frequent than duplications, an interstitial duplication at 9p24.3, encompassing the genes *DOCK8* and *KANK1*, has been recently associated with multiple neurodevelopmental disorders, including schizophrenia (SZ), bipolar disease, autism spectrum disorders (ASD), attention deficit hyperactivity disorder (ADHD), and depression, all of them conditions entailing problems with language (Glessner et al., 2017).

In this paper, we report on a boy with a partial duplication of *DOCK8* resulting from a de novo microduplication in 9p24.3 and provide a detailed characterization of his language (dis)abilities. A growing body of evidence points to an association of *DOCK8* deletions, duplications, and translocations interrupting its functionality, with intellectual disability and developmental delay (Griggs et al., 2008; Krgovic et al., 2018). According to DECIPHER (https://decipher.sanger.ac.uk/), nearly 60% of the 143 CNVs involving *DOCK8* are losses and about 40% are gains. *DOCK8* losses are associated with intellectual disability, global developmental delay, abnormal facial features, and microcephaly, whereas *DOCK8* gains are mostly associated with intellectual disability, global developmental delay, ASD, clinodactyly, and delayed speech and language development. However, these CNVs usually involve other genes besides *DOCK8* and/or are found in patients with other chromosomal abnormalities. Accordingly, we expect that smaller CNVs affecting to *DOCK8* only, like the one found in our patient, help to achieve more robust genotype-to-phenotype correlations.

*DOCK8* plays an important role in modulating the immune response and mutations in the gene are known to cause immunodeficiency disease (Biggs et al., 2017; Broides et al., 2017; Kearney et al., 2017). Because the gene is also expressed in the fetal and adult brains (Griggs et al., 2008), it is expected to play a key role as well in brain development and function. Accordingly, in mice the Dock8 protein contributes to Schwann cell precursor migration during embryonic development of the peripheral nervous system (Miyamoto et al., 2016). *DOCK8* is upregulated in neuronally differentiated cells after the over-expression of MECP2_e1, which is the etiologically relevant variant of the MECP2 protein for Rett syndrome (Orlic-Milacic et al., 2014). *DOCK8* is a candidate for ASD too. Two de novo loss-of-function variants of *DOCK8* have been found in probands from the Autism Sequencing Consortium and the Autism Clinical and Genetic Resources in China (ACGC) cohort (De Rubeis et al., 2014; Wang et al., 2016). The 9p24.3 region overlaps with linkage regions found in large ASD extended pedigrees (Allen-Brady et al., 2009; Coon et al., 2010). Nonetheless, little is known about the causes of the neuropsychiatric features and neurodevelopmental anomalies associated to mutations of *DOCK8* or to CNVs encompassing the gene. This is particularly true of the language deficits that are commonly observed in patients, among other reasons, because we still lack a detailed account of the language problems caused by *DOCK8* mutations or dosage alterations. In this paper, we build on our current knowledge of the genetic aspects of language development and language evolution, as well as on the expression pattern of selected genes in the blood of our proband, for advancing a hypothesis of the molecular causes of language dysfunction associated to *DOCK8* alterations.

## MATERIAL AND METHODS

### Linguistic, cognitive, and behavioral assessment

The global developmental profile of the proband was defined with the Spanish versions of the Wechsler Intelligence Scale for Children, Fifth Edition (WISC-V) (Hernández et al., 2015) and the Inventory for Client and Agency Planning (ICAP) (Montero, 1996).

#### Wechsler Intelligence Scale for Children (WISC-V)

The WISC-V was used for testing multiple cognitive abilities of the child. It consists of 10 primary subtests and 5 secondary subtests, which are combined to generate different composite score indices. The five primary composite score indices are: Verbal Comprehension (VCI), Visual Spatial Index (VSI), Fluid Reasoning Index (FRI), Working Memory Index (WMI), and Processing Speed Index (PSI). The VCI measures verbal reasoning, understanding, concept formation, and crystallized intelligence. The VSI measures nonverbal reasoning and concept formation, visual perception and organization, visual-motor coordination, and the child’s ability to analyze and synthesize abstract information, and to distinguish figure-ground in visual stimuli. The FRI evaluates quantitative reasoning, classification and spatial abilities, knowledge of part to whole relationships, as well as fluid reasoning abilities. The WMI assesses the child’s ability to sustain auditory attention, concentrate, and exert mental control. Lastly, the PSI determines the speed and accuracy of the child when processing information.

#### The Inventory for Client and Agency Planning (ICAP)

The ICAP was used for evaluating the subject’s functional abilities and maladaptive behaviors in the following general areas: motor skills, social and communication skills, personal living skills, and community living skills. The ICAP measures the frequency and severity of 8 types of behavioral disturbances, which are organized in 3 subscales: asocial maladaptive behavior (i.e. uncooperative behavior and socially offensive behavior), internalized maladaptive behavior (i.e. withdrawn or inattentive behavior, unusual or repetitive habits, and self-harm), and externalized maladaptive behavior (i.e. disruptive behavior, destructive to property, and hurtful to others). Behavior is rated as normal or abnormal, whereas behavioral problems are subsequently rated as marginally serious, moderately serious, serious, or very serious.

Regarding the language profile of our proband, his abilities were evaluated in depth with the Spanish versions of the Clinical Evaluation of Language Fundamentals, Fourth Edition (CELF-4) (Semel et al., 2006), the Peabody Picture Vocabulary Test, Third Edition (PPVT-3) (Dunn et al. 2006), the Test de Comprensión de Estructuras Gramaticales [Test for evaluating the comprehension of grammatical structures] (CEG) (Mendoza et al., 2005), and the Registro Fonológico Inducido [Triggered Phonological Register] (Juárez and Monfort, 1998).

#### *Clinical Evaluation of Language Fundamentals* (CELF-4)

This test was used to evaluate the language (dis)abilities of the child and to gain a general view of his strengths and deficits during communication, and of the language domains that were more severely impaired. For subjects above 9 years old, like our proband, the CELF-4 encompasses 16 subtests, which are organized in four levels. Level 1 is aimed to identify whether or not the child suffers from a language disorder (expressed by the Core Language Score, CLS). Level 1 comprises the following 5 subtests: 1) Concepts & Following Directions (C&FD), in which the subject is expected to point to images of objects in response to oral commands, and which is aimed to evaluate her ability to interpret spoken directions, remember names and features of objects, and identify them among other pictured objects; 2) Recalling Sentences (RS), in which the subject is expected to imitate different target sentences and which is aimed to assess his ability to listen and repeat sentences in an accurate way; 3) Formulated Sentences (FS), in which the subject is expected to formulate full sentences about different visual stimuli that are presented using target words or phrases, and which is aimed to evaluate his ability to create well-formed and congruent spoken sentences relying on given words and the contextual information provided by illustrations; 4) Word Classes 2-Receptive (WC 2-R), in which the subject is expected to point to the two pictured objects that are most related among a set of different objects, and which is aimed to evaluate his ability to understand logical relationships in the meanings of associated words; and 5) Word Classes 2-Expressive (WC 2-E), in which the subject is expected to describe the relationship that holds between the two objects chosen in subtest WC 2-R, and which is aimed to assess his ability to explain logical relationships in the meanings of associated words. At this Level 1 the WC 2-R and the WC 2-E scores are usually merged in a Word Classes Total score (WC-T).

Level 2 is aimed to identify the nature of the disorder through 4 index scores, resulting from different subtests. The Receptive Language Index (RLI) measures listening and auditory comprehension (Receptive Language Index, RLI). It is calculated from the scores obtained in the C&FD and the WC 2-R subtests. The Expressive Language Index (ELI) measures the expressive language skills and is obtained from the scores obtained in the RS, FS, and WC 2-E subtests. The Language Content Index (LCI) measures the subject’s ability to understand and produce linguistic meaning by means of a set of subtests assessing the proband’s lexical and morphosyntactic knowledge along with his pragmatic skills. It results from the WC-T score, plus the scores obtained in two other specific subtests: Words Definitions (WD) and Understanding Spoken Paragraphs (USP). In the WD the subject is expected to provide a definition of several target words, that are orally presented, followed by an introductory sentence that includes the word. This subtest is aimed to evaluate the subject’s expressive vocabulary, and particularly, his ability to analyze words for their meaning features, define words by referring to class relationships and shared meanings, and describe meanings that are unique to the reference or instance. In the USP the subject is expected to answer to different questions about a paragraph presented orally by the examiner. The questions target the paragraph’s main idea, some of its details, and its sequencing, but also different kind of inferential and predictive information. This subtest is aimed to assess the subject’s ability for sustained attention and focusing, but also to understand oral texts, and respond to questions about the contents of the given information. Finally, the Language Memory Index (LMI) measures the subject’s ability to apply working memory to linguistic content and structure. It is calculated from the scores obtained in the C&FD, RS, and FS subtests.

Level 3 is aimed to assess underlying behaviors of clinical significance that may account for the attested language problems. It makes use of 3 specific subtests: 1) Phonological Awareness (PA), in which the subject is expected to identify, segment, blend, or delete sounds and syllables, and which evaluates his knowledge of the sound structure and rules of his language, and his ability to manipulate the sound units making up words and sentences; 2) Word Associations (WA), in which the subject is asked to recall words of specific semantic categories while being timed, and which is aimed to evaluate his semantic category learning and knowledge, in turn a measure of the development and adequacy of his vocabulary knowledge and use in receptive and expressive tasks; 3) Rapid Automatic Naming (RAN), in which the subject is asked to name familiar colors, shapes, and shape-color combinations while being timed, and which is aimed to evaluate his ability to produce automatic speech. Level 3 also includes a Working Memory Index (WMI), which assesses working memory (important for attention, concentration, and recall) through 2 specific subtests: Number Repetition 2 Total (NR 2-T) and Familiar Sequences 2 (FS 2). In the NR2-T the subject is asked to repeat series of numbers forward and then backwards, with the aim of evaluating his abilities to repeat random number sequences. In the FS the subject is asked to name the days of the week, count backward, or recall other kind of information while being timed, with the aim of evaluating his ability to quickly sequence auditory and verbal information, but also his attention, concentration, processing speed, and auditory/verbal working memory.

Finally, Level 4 is aimed to evaluate language and communication in use, by eliciting information from parents, teachers, or the subject himself about the subject’s social and communication skills. This level encompasses 2 tasks: Pragmatics Profile (PP) and Observational Rating Scale (ORS). In the PP the examiner elicits information from a parent or teacher about the subject’s social language skills. This task is aimed to determine his overall pragmatic development. In the ORS a parent, a teacher, and the subject each rate the subject’s interaction and communication skills.

#### *The Peabody Picture Vocabulary Test* (PPVT-3)

This test was used to evaluate in more detail the receptive abilities of the proband. The PPVT-3 is a performance test that assesses vocabulary acquisition via several tasks in which the child is asked to point to one of four color pictures on a page after hearing a word. The test is designed for subjects from 2.5 years to 90 years and it can also be used for a rapid screening of difficulties with language or abnormal verbal attitudes.

#### Test de Comprensión de Estructuras Gramaticales (CEG)

This test, which is based on the Test for Reception of Grammar (TROG-2) (Bishop, 2003), was also used to gain a more accurate view of the receptive abilities of the proband. Similarly to the PPVT-3, the CEG does not demand any verbal response from the child, which is asked to point to one of four pictures (one correct picture and three distractors) after listening to the target sentences. Each grammatical structure is tested via 4 different items. 20 different grammatical constructions are assessed (see Supplemental file 1). The resulting scores enable the quantification of the developmental delay of the proband, as well as the identification of the grammatical aspects that are more problematic to him.

#### Registro Fonológico Inducido

This tests assesses phonological and articulatory abilities of Spanish-speaking children between 3 and 7 years old by means of elicited word production and repetition. Word production is induced by showing the child different pictures that she has to name. If the child is unable to produce the name, the speech-language therapist will provide her with the correct name and will ask the child to repeat it. The test comprises every possible sound of the Spanish language, so that it can be used to detect dyslalia as well as phonological impairment.

#### Natural speech sample analysis

Additionally, we assessed the proband’s language in use and overall communication skills through the systematic analysis of a 15-minute sample of the child’s talk while he was playing and chatting about familiar topics (family and school) with his mother and his speech language therapist (MSJR, one of the authors of the paper). The conversation was video-recorded and then transcribed and coded using CHAT (Codes for the Human Analysis of Transcripts) software, a tool of the CHILDES Project (MacWhinney, 2000). CHAT codes the speech production phenomena on the main lines of the transcript, whereas the phonological, morphological, syntactic and pragmatic phenomena are coded on the dependent lines. Information about prosody and communicative gestures and gaze is included within square brackets in the main lines. Pauses are also coded, with an estimation of duration (see https://talkbank.org/manuals/CHAT.pdf for details). We then proceeded to evaluate the most relevant aspects of language in use, which entails assessing every linguistic phenomenon from a pragmatic perspective (Verschueren, 1999). Hence, we first tagged the child’s errors, with an indication of their nature (phonological, morphological, syntactic, lexical semantic). Afterwards, we determined their impact on communication and labelled them according to PREP-CORP (PRagmatic Evaluation Protocol for the analysis of oral CORPpora), which has been previously used for obtaining the pragmatic profile of other developmental disorders (Fernández-Urquiza et al., 2015; Fernández-Urquiza et al., 2016; Shiro et al., 2016; Diez-Itza et al., 2018). Lastly, a global description of the pragmatic linguistic profile of the proband was built up by automatically computing the number of occurrences of each label by means of CLAN (Computerized Language Analysis) (Conti-Ramsden, 1996).

### Cytogenetic and molecular analyses

#### Karyotype analysis

Peripheral venous blood lymphocytes were grown following standard protocols and collected after 72 hours. A moderate resolution G-banding (550 bands) karyotyping by trypsin (Gibco 1x trypsin^®^ and Leishmann stain) was subsequently performed. Microscopic analysis was conducted with a Nikon^®^ eclipse 50i optical microscope and the IKAROS Karyotyping System (MetaSystem^®^ software).

DNA from the patient and his parents was extracted from 100 μl of EDTA-anticoagulated whole blood using MagNA Pure (Roche Diagnostics, West Sussex, UK) and used for subsequent analyses.

#### Fragile X syndrome analysis

CGG expansions affecting the gene *FMR1* (the main determinant factor for X-fragile syndrome) were analyzed in the proband according to standard protocols. Polymerase chain reaction (PCR) of the fragile site was performed with specific primers for the fragile region of the *FMR1 locus* and the trinucleotide repeat size of the resulting fragments was evaluated by electrophoresis in agarose gel.

#### Multiplex ligation-dependent probe amplification (MLPA)

MLPA was conducted to detect abnormal copy-number variations (CNVs) in subtelomeric regions of the chromosomes, as well as frequent interstitial CNVs. MLPA consists on the amplification of different probes using a single PCR primer pair. Each probe detects a specific subtelomeric DNA sequence. Two different kits from MRC-Holland were used: SALSA^®^ MLPA^®^ probemix P036-E1 and SALSA^®^ MLPA^®^ probemix P070-B2 Human Telomere-5. SALSA^®^ MLPA^®^ probemix P036-E1 contains 46 MLPA probes with amplification products between 130 and 483 nt: 2 probes for each chromosome. 41 probes are located in subtelomeric regions. No probes are present for the subtelomeric regions of the 5 acrocentric chromosomes (13, 14, 15, 21, 22). For these, an extra probe is included detecting the q arm, close to the centromere. The subtelomeric probes for the X and Y chromosome are identical as they detect sequences in the pseudoautosomal regions (PAR1 and PAR2) which are identical in chromosome X and Y. The following genes are detected by the probes included in this probemix: *TNFRSF4, SH3BP5L, ACP1, CAPN10, CHL1, BDH1, PIGG, TRIML2, PDCD6, GNB2L1*, *IRF4*, *PSMB1*, *ADAP1*, *VIPR2*, *FBXO25*, *ZC3H3*, *DMRT1*, *EHMT1*, *DIP2C*, *PAOX, RIC8A, NCAPD3, SLC6A12, ZNF10, PSPC1, F7, CCNB1IP1, MTA1, MKRN3, ALDH1A3, POLR3K, GAS8, RPH3AL, TBCD, USP14, RBFA, CDC34, CHMP2A, SOX12, OPRL1, RBM11, PRMT2, BID, RABL2B, SHOX* and *VAMP7*, respectively.

Regarding SALSA^®^ MLPA^®^ probemix P070-B2 Human Telomere-5 it also contains 46 MLPA probes with amplification products between 132 and 484 nt: 2 probes for each chromosome. As with the former probemix, no probes are present for the subtelomeric regions of the 5 acrocentric chromosomes (13, 14, 15, 21, 22). For these, an extra probe is included detecting the q arm, close to the centromere. Likewise, the subtelomeric probes for the X and Y chromosome are also identical as they detect sequences in the pseudoautosomal regions (PAR1 and PAR2) which are identical in chromosome X and Y. The following genes are detected by the probes included in this probemix: *TNFRSF18, SH3BP5L, ACP1, ATG4B, CHL1, KIAA0226, PIGG, FRG1, CCDC127, GNB2L1, IRF4, TBP, SUN1, VIPR2, FBXO25, RECQL4, DOCK8, EHMT1, ZMYND11, ECHS1, BET1L, IGSF9B, JARID1A, ZNF10, PSPC1, CDC16, PARP2, MTA1*, *NDN*, *TM2D3*, *DECR2*, *GAS8*, *RPH3AL*, *SECTM1*, *THOC1*, *CTDP1*, *PPAP2C*, *CHMP2A*, *ZCCHC3*, *UCKL1*, *HSPA13*, *S100B*, *IL17RA*, *ARSA*, *SHOX* and *VAMP7*, respectively.

The PCR products were analyzed by capillary electrophoresis in an automatic sequencer Hitachi 3500 and further analyzed with the Coffalyser V 1.0 software from MRC-Holland.

#### Microarrays for whole-genome CNVs search and chromosome aberrations analysis

The DNA from the patient and his parents was hybridized on a CGH platform (Agilent Technologies). The derivative log ratio spread (DLRS) value was 0.11. The platform included 60.000 probes. Data were analyzed with the Agilent Genomic Workbench 7.0, and the ADM-2 algorithm (threshold = 6.0; aberrant regions had more than 5 consecutive probes).

In order to determine the genes that could be differentially expressed in the proband (DEGs) compared to his healthy parents, microarray analyses of blood samples from the three of them were performed. Total RNA was extracted with the PAXgene Blood RNA Kit IVD (Cat No./ID: 762164). RNA quality and integrity were confirmed with a Bioanalyzer RNA 6000 Nano. All samples had RNA integrity number (RIN) values above 9. An Affymetrix^®^ Scanner 3000 7G was then used for analyzing transcriptome changes. The resulting raw data were processed with the Affymetrix^®^ GeneChip^®^ Command Console^®^ 2.0 program. Next, *.CEL files were checked to certify the RNA integrity and the suitability of the labeling and the hybridization processes. Finally, the raw data from the different arrays were normalized with the SST (Signal Space Transformation)-RMA (Robust Microarray Analysis) tool (Irizarry et al., 2003). Normalized data (*.CHP files) were subsequently used to search for DEGs in the proband compared to his parents. Statistical analyses were conducted with the LIMMA (Linear Models for Microarray Analysis) package of BioConductor, using the TAC 4.0 software.

## RESULTS

### Clinical History

The proband is a boy born after 9 months of gestation by caesarean section. At birth, his weight was 3450 g (65^th^ percentile), his height was 50 cm (54^th^ percentile), and his occipitofrontal circumference was 37 cm (98^th^ percentile). The Apgar scores were 9 (at 1’) and 9 (at 5’). At the delivery, no signs of disease were observed and the newborn was fed with both maternal and artificial milk. Cranial magnetic resonance imaging (MRI), performed at 4 years of age, yielded no pathological results. The child has suffered from recurrent sinusitis, resulting in frequent nose infections and headaches. At the moment of our evaluation, he was 11 years and 6 months old. He exhibited mild dysmorphic facial features, particularly, hypertelorism and mild retrognathia.

### Language and cognitive development

Early developmental milestones were achieved normally by the child, including sphincter control at the age of 2 years. Cognitive anomalies were first reported by the teachers of the nursery school when the proband was around 2 years old. At the age of 3 years, language delay was evident. According to his parents, the child’s language was gibberish, with poor articulation, and he usually aided himself with body and hand gestures to communicate better. The child received early childhood intervention from the age of 4 years to the age of 6 years, and speech therapy from the age of 3 years to present. At the age of 6 years and 1 month, the child was evaluated by the hospital’s neuropsychologist, because of his language delay and learning difficulties. His language skills were poor. He uttered pseudosentences consisting of 3-4 content words with no functional words, like articles. Expressive language was more impaired than comprehension. The child scored low in the Leiter International Performance Scale, 3rd Edition (Leiter-3), a test aimed to evaluate nonverbal cognitive, attentional, and neuropsychological abilities in typical and atypical populations (Roid et al., 2011). The resulting intellectual quotient (IQ) was 66, with an estimated mental age of 4 years (more than 2 years below his chronological age). He was diagnosed of specific language impairment and developmental delay. Motor delay was attested too.

At the age of 9 years, our proband was evaluated by a child psychologist. The boy scored abnormally high in the Spanish version of the Conners Comprehensive Behavior Rating Scales (Conners CBRS) (Conners, 2008), as well as abnormally high (1^st^ percentile) in the Escalas para la Evaluación del Trastorno por Déficit de Atención con Hiperactividad [Scales for the Evaluation of ADHD] (EDAH) (Farré and Narbona, 2013), this being suggestive of the presence of ADHD. Likewise, he scored very low in the Spanish version of the Differences Perception Test (CARAS-R) (Thurstone and Yela, 2012), which suggested that he was unable to judge properly about similarities and differences and paid reduced attention to details, because of his reduced visuo-perceptive abilities. He also scored very low in the impulsivity control index, which pointed to a high impulsivity and lack of inhibitory control. In the Spanish version of the Behavior Assessment System for Children (BASC-2) (Reynolds and Kamphaus, 2004), which evaluates different aspects of the child’s behavior, our proband scored abnormally high in the subtests assessing the presence of aggressiveness, hyperactivity, attention deficit, and depression. Finally, regarding his intellectual development, the boy obtained 77 points (6^th^ percentile) in the Spanish version of the Reynolds Intellectual Assessment Scales (RIAS) (Reynolds and Kamphaus, 2013). He obtained 82 points (11^th^ percentile) in the Verbal Index of the test, 77 points (6^th^ percentile) in the Non Verbal Index, and 80 points (9^th^ percentile) in the Memory Index. Overall, the obtained scores pointed to a mild-to-moderate cognitive impairment impacting on his functional abilities. Similarly, the IQ of our proband according to the Spanish version of the Cattell intelligence test (Cattell and Cattell, 1994) was 65 (4^th^ percentile), which was indicative of low-to-moderate intellectual disability.

At the age of 11 years the proband was again evaluated by the school’s psychologist, who reported cognitive delay, difficulties for recalling information, problems for understanding, poor reasoning, and problems with spatio-temporal perception. He also exhibited low working memory abilities. In the language domain, articulatory problems were observed, in part resulting from an orofacial hypotonia, as well as expressive and comprehensive deficits in the oral and written domains. Although our subject has learnt to read and write, he makes frequent mistakes. He also exhibited behavioral disturbances, including low tolerance to frustration, variable mood, frequent distractions, and difficulties for interacting with some of his peers (being instead attentive and kind with adults). Minor coordination problems in the motor domain were confirmed. However, they did not preclude him from mastering the needed skills for skiing, biking, or skating. At that moment, he was attending a normal school, although he was aided by an assistant teacher several hours a week.

When the child was 11 years and 6 months old, we assessed in detail his global development with the Spanish versions of the Wechsler Intelligence Scale for Children, Fifth Edition (WISC-V) and the Inventory for Client and Agency Planning (ICAP). In the WISC-IV the child scored low in most of the subtests (Table 1), this resulting in a IQ of only 54. As also shown in Table 1, verbal comprehension, fluid reasoning, working memory, and processing speed are significantly impaired, being the visual spatial abilities the most preserved domain. Regarding the ICAP, which was administered to assess the child’s adaptive behavior, the obtained scores were indicative of a delay of 5 years and 3 months in his global development, being social and communication skills the most impaired domain, whereas community living skills were relatively spared (Figure 1A; see Supplemental file 1 for details).

**Table 1.** Summary table of the scores obtained by our proband in the WISC-V test. Subtests used to derive the IQ are bolded. Secondary subtests are in parentheses. Abbreviations: SEM: standard error of measurement; y: years; m: months.

| Scale | Subtest Name | Total Raw Score | Scaled Score | Percentile Rank | Age Equivalent | SEM | Total Scaled Score | Composite Score |
| --- | --- | --- | --- | --- | --- | --- | --- | --- |
| Verbal Comprehension | **Similarities** | 15 | 3 | 1 | <6y 2m | 1.04 | 4 | 55 |
|  | **Vocabulary** | 12 | 1 | 0.1 | <6y 2m | 1.27 |  |  |
|  | (Information) | 8 | 2 | 0.4 | <6y 2m | 1.08 |  |  |
|  | (Comprehension) | 12 | 3 | 1 | 6y 2m | 1.37 |  |  |
| Visual Spatial | **Block Design** | 14 | 4 | 2 | <6y 2m | 1.41 | 11 | 75 |
|  | Visual Puzzles | 12 | 7 | 16 | 7y 10m | 0.99 |  |  |
| Fluid Reasoning | **Matrix Reasoning** | 5 | 1 | 0.1 | <6y 2m | 1.04 | 8 | 67 |
|  | **Figure Weights** | 17 | 7 | 16 | 8y 6m | 0.73 |  |  |
|  | (Arithmetic) | 10 | 3 | 1 | 6y 6m | 0.85 |  |  |
| Working Memory | **Digit Span** | 18 | 5 | 5 | 6y 10m | 0.90 | 8 | 67 |
|  | Picture Span | 11 | 3 | 1 | <6y 2m | 1.34 |  |  |
|  | (Letter-Number Seq.) | 5 | 1 | 0.1 | <6y 2m | 1.20 |  |  |
| Processing Speed | **Coding** | 15 | 1 | 0.1 | <8y 2m | 1.31 | 5 | 56 |
|  | Symbol Search | 13 | 4 | 2 | <8y 2m | 1.37 |  |  |
|  | (Cancellation) | 65 | 9 | 37 | 10y 6m | 1.20 |  |  |
| **Total Scale** |  |  |  |  |  |  | **22** | **54** |

**Figure 1.**
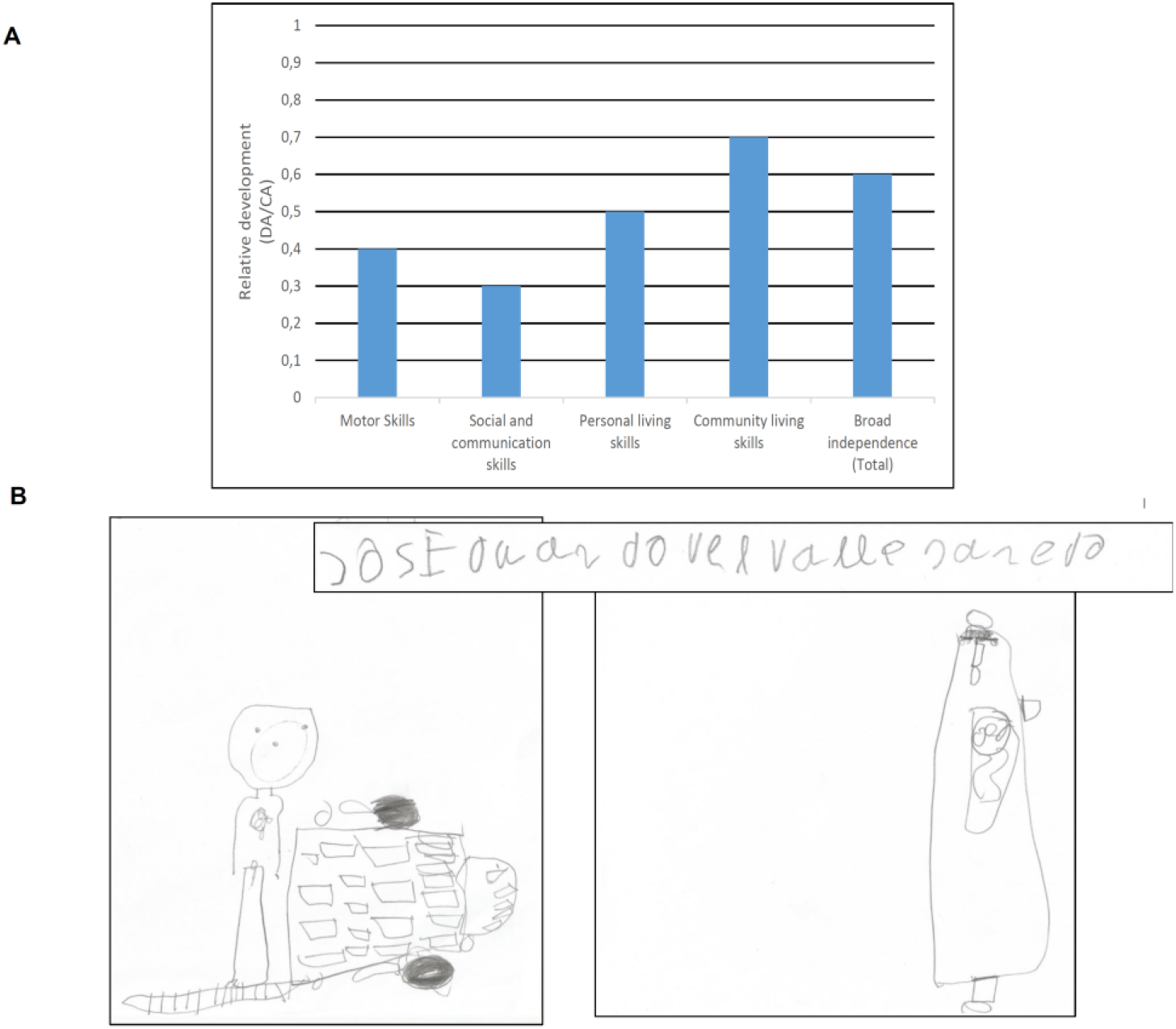
Distinctive clinical features of the proband. A. Developmental profile of the proband at age 11 years and 6 months according to the Inventory for Client and Agency Planning (ICAP). Abbreviations: DA, developmental age; CA, chronological age. B. Samples of the child’s writing and drawing abilities. His handwritten name is displayed above. Below two different drawings are shown: the proband’s sister leaving for a plane trip (left) and a motorcycle (right). The size of the drawings correlate with their emotional contents.

Because of the significant impairment reported by the WISC-V, we assessed in detail the linguistic abilities of the proband at the age of 11 years and 6 months, using different tests aimed to evaluate his expressive and comprehensive abilities, as well as how they were put into use for communicating.

Table 2 summarizes the scores obtained by our proband in the subtests comprising the CELF-4, which is aimed to provide a detailed overview of the language deficits and strengths of the subject. The Core Language Score of the CELF-4 was indicative of a moderate language disorder. The most affected domains were Working Memory and Language Memory, which were severely impaired. Language Content, Receptive Language and Expressive Language were moderately affected. Interestingly, as also shown in Table 2, in some of the subtests our proband scored like neurotypical children, significantly in tasks evaluating semantic knowledge. Conversely, he obtained the lowest scores in tasks assessing his ability to understand and compose sentences, and to retrieve and process verbal information. Regarding the subtests aimed to establish the nature of the underlying cognitive processes plausibly accounting for these problems (Level 3 of the CELF-4), he scored atypically (in terms of time of execution and performed errors) in the tasks assessing his ability to produce automatic speech (Rapid Automatic Naming), and scored very low in the Working Memory Index (Table 2). On the contrary, he scored only slightly below his age-matched peers in the tasks evaluating his phonological competence (Phonological Awareness) and his semantic category knowledge (Word Associations). Lastly, regarding his pragmatic abilities (Level 4 of the CELF-4), our proband exhibited problems for properly putting his language knowledge into use. The Pragmatic Profile suggested that he was performing below his chronological age. The checklist in the Observational Rating Scale component corroborated the findings at the other levels of the test, pointing to widespread problems in all the assessed domains: language comprehension, language production, and literacy. The proband was reported to have difficulties for finding the correct or needed words, and for describing things, expressing what he meant, and describing events in a coherent fashion. Likewise, he was reported to have problems with sustained attention and with following and recalling oral instructions. On the contrary, he was able to follow the other’s gaze and paid attention to non-verbal, gestural and corporal language during conversational exchanges. Moreover, he seemingly had a good command of the principles regulating conversation (for instance, he knows how to ask for clarifications and for aid when he does not understand something). Lastly, this part of the test also corroborated his generalized problems with reading and writing. He experiences difficulties for understanding written texts and extracting the main theme. Likewise, he exhibited difficulties for writing, particularly with the shape of the letters, and for drawing, although his drawing abilities are in line with his cognitive deficits and language problems (Figure 1B).

**Table 2.**
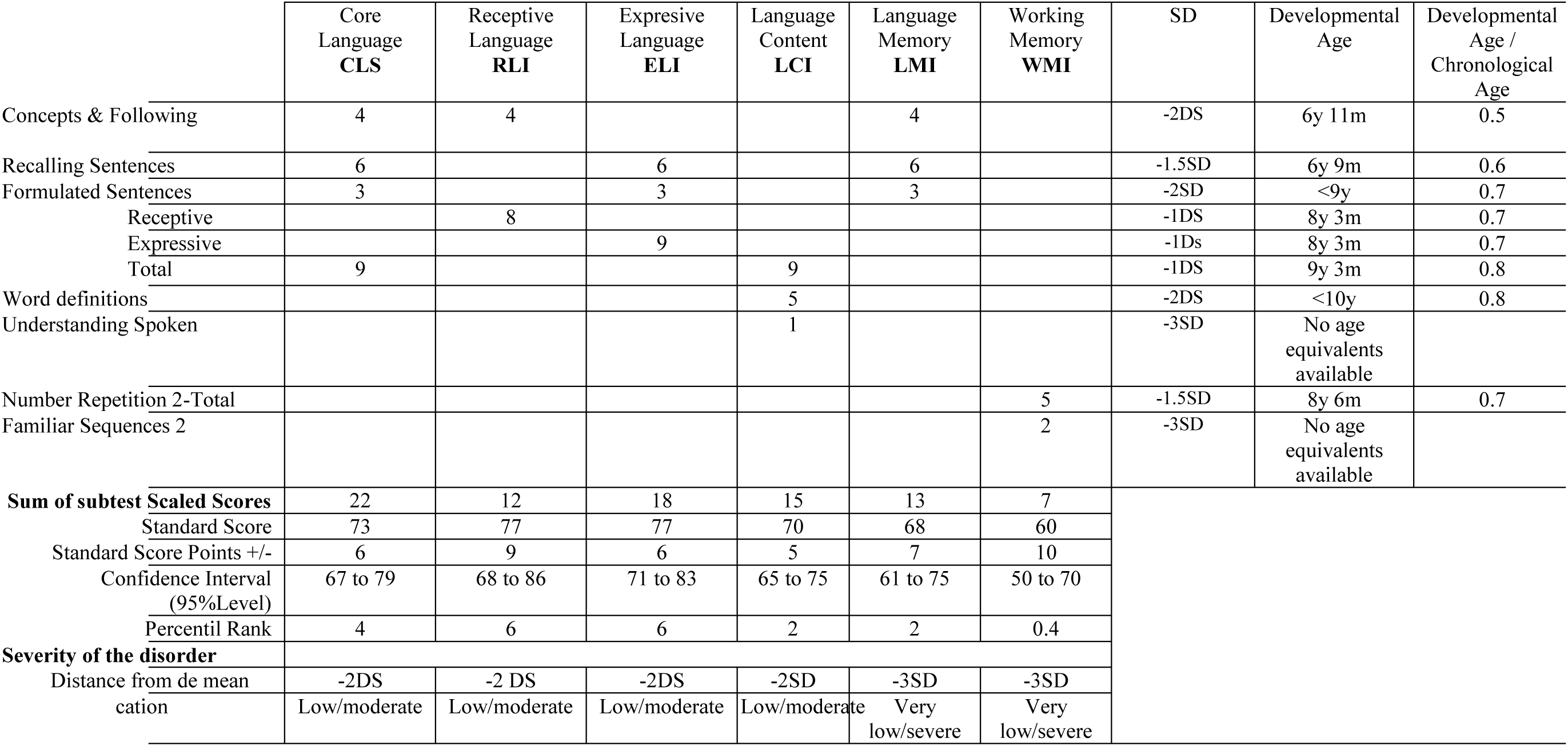
Summary table of the scores obtained by our proband in the CELF-4 test. Abbreviations: SD: standard deviation; y: years; m: months.

We assessed in more detail aspects of the phonetic, semantic, and grammatical profile of our proband, in order to refine our view of his deficits and strengths in the language domain. Articulatory problems were evaluated with the Registro Fonológico Inducido, which pointed out that he has problems mostly with liquids. Accordingly, he regularly assimilates [ɾ] to [l], like in [ˈflesa], instead of [ˈfɾesa] ‘strawberry’, or [ˈbluja], instead of [ˈbɾuja] ‘witch’. Polysyllabic words were also problematic for him, as in [pɾeˈðɾjoðiko] instead of [peˈɾjoðiko] ‘newspaper’ (see Supplemental file 1 for details). Semantic problems were assessed in more depth with the Peabody Picture Vocabulary Test (PPVT-3). The resulting scores pointed to a delay of 2 years in his receptive vocabulary (see Supplemental file 1 for details). Finally, we used the Test de Comprensión de Estructuras Gramaticales (CEG) for refining our characterization of the proband’s grammatical knowledge. The global score obtained by the child in this test was 62 (10^th^ percentile). This low performance was mostly caused by generalized problems for correctly computing marked word order (like in topicalized or cleft sentences), anaphora (particularly, in complex sentences with relative pronouns) and clitics. On the contrary, sentence length did not have any significant impact on his performance (see Supplemental file 1 for details).

Finally, we examined in even more depth the child’s language in use and his communication skills through the analysis of several naturally-occurring interactions with his mother and his speech therapist (see Supplemental file 2 for details). We confirmed that the proband’s speech exhibits frequent (but non-systematic) substitutions of specific consonants and vowels (e.g. [r] > [d], [ɾ] > [d], [d] > [ɾ], [s] > [θ], [ɾ] – [r] > [l], [e] > [a], [e] > [i], [o] > [a]), as well as recurrent simplification of consonant clusters (e.g. [taɾˈxeta] > [taˈxeta] ‘card’, [pɾiˈmeɾo] > [piˈmeɾo] ‘first’, [ˈeɾnja] > [ˈennja] ‘hernia’, [tɾaβaˈxaɾ] > [taβaˈxaɾ] ‘work’, [ˈgusta] > [ˈguta] ‘I like’, [ˈgɾita] > [ˈɾita] ‘he shouts’, 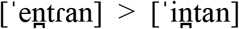 ‘they go into’). We also found occasional diphthong simplifications (e.g. [we] > [e]), sound assimilations ([deθiˈmales] > [diθiˈmales] ‘decimals’; [ˈposteɾ] > [ˈpotteɾ] ‘poster’), and metatheses or sound inversions (e.g. [aðxeˈtiβos] > [axeðˈtiβos] ‘adjectives’). Speech prosody and fluency were significantly impaired due to the articulatory difficulties experienced by the child, which hinders his speech intelligibility.

Concerning language structure and complexity, the mean length of utterance in words (MLUw) was 4.2, which is within the range of children aged between 4 years and 6 months, and 5 years (Rice et al., 2010), and thus remarkably below his chronological age. The child produced morphosyntactic errors such as substitutions of inflectional morphemes (dicen > dice ‘they say’, as in *y todos dice ‘and all of them say’), substitution of conjunctions (que > a ‘that’, as in *tenemos a poner ‘we have to put’), and omissions of prepositions (de > ø ‘of’ as in *qué color soy ‘(of) which color I am’).

Finally, we observed that the child usually avoided the indirect or reported speech, which is more complex syntactically. Instead, when requested to retell a story with several characters, he represented the different perspectives by dramatizing each character by means of direct reported speech, as is commonly observed in people with Williams Syndrome (Losh et al., 2000; Reilly et al., 2004; Gonçalves et al., 2010; Shiro et al., 2016).

Despite these problems with some structural components of language, the boy’s interactive pragmatic skills seemed to be quite preserved. Accordingly, he had mastered most of the conversational abilities, including turn-taking and repair strategies (Sacks et al., 1974). For instance, he made an adequate communicative use of gaze when taking the floor, but also when ceding it to the other participants. Likewise, in order to compensate his expressive difficulties, he might explicitly request for help and make visual contact with his interlocutor. Additionally, he used fillers, word repetitions, and reformulations to keep the turn and extend the time of elocution, in order to gain some extra seconds to build up his utterances. Likewise, his echolalic replies reflected his willingness to collaborate with the interlocutor during verbal interactions despite his comprehension problems. Interestingly, the child made extensive use of interjections along with facial expressions as a sort of “inference-launching mechanism” that seemingly allowed him to communicate implicit propositional contents avoiding at the very same time his expressive difficulties. That is to say, the intentional transgression of the conversational maxims of manner (‘be as clear, as brief, and as orderly as you can in what you say, and avoid obscurity and ambiguity’) and quantity (‘be as informative as possible and give as much information as needed, but no more’) resulted in successful implicit communication. However, pathological transgressions of the maxims of quality (‘be truthful and do not give information that is false or that is not supported by evidence’) and relation (‘be relevant and say things that are pertinent to the discussion’) were also observed. For example, when asked simple questions about the characteristics of a pineapple, he would say that it is not yellow, but triangular and with seeds. This might result from an aberrant reasoning and/or lack of awareness of the world of reference shared by his interlocutors, which might explain as well his difficulties to understand even the dynamics of simple games.

### Cytogenetic and molecular analyses

Routine cytogenetic and molecular analyses of the proband were performed when he was 6 years and 6 months old. The karyotype was normal, although the analysis did not discard the presence of minor chromosomal aberrations. PCR analysis of the *FMR1* fragile site was normal (28 repetitions). MLPAs of subtelomeric regions detected an increase of gene dosage in 9p24.3 involving *DOCK8* (P070-B2 Human Telomere-5 probemix). A comparative genomic hybridization array (array-CGH) was performed a month later, when the proband was 6 years and 7 months old. The arrays confirmed the presence of a duplication of nearly 0.2 Mb in the 9p24.3 region (arr[hg19] 9p24.3(266,045-459,076)x3), predicted to be probably benign, and affecting the gene *DOCK8* (figures 2A and 2B). However, it also uncovered a microdeletion of 0.5 Mb in the 8p23.1 region (arr[hg19] 8p23.1(7,169,490-7,752,586)x1), predicted to be benign. This microdeletion in chromosome 8, encompasses 19 protein-coding genes: 12 genes coding for defensins, 2 genes coding for proline rich domain proteins, 2 genes coding for sperm associated antigen proteins, 2 genes coding for ubiquitin peptidase-like proteins, and 1 gene coding for a zinc finger protein (figures 2C and 2D). Both alterations occurred de novo in the proband.

**Figure 2.**
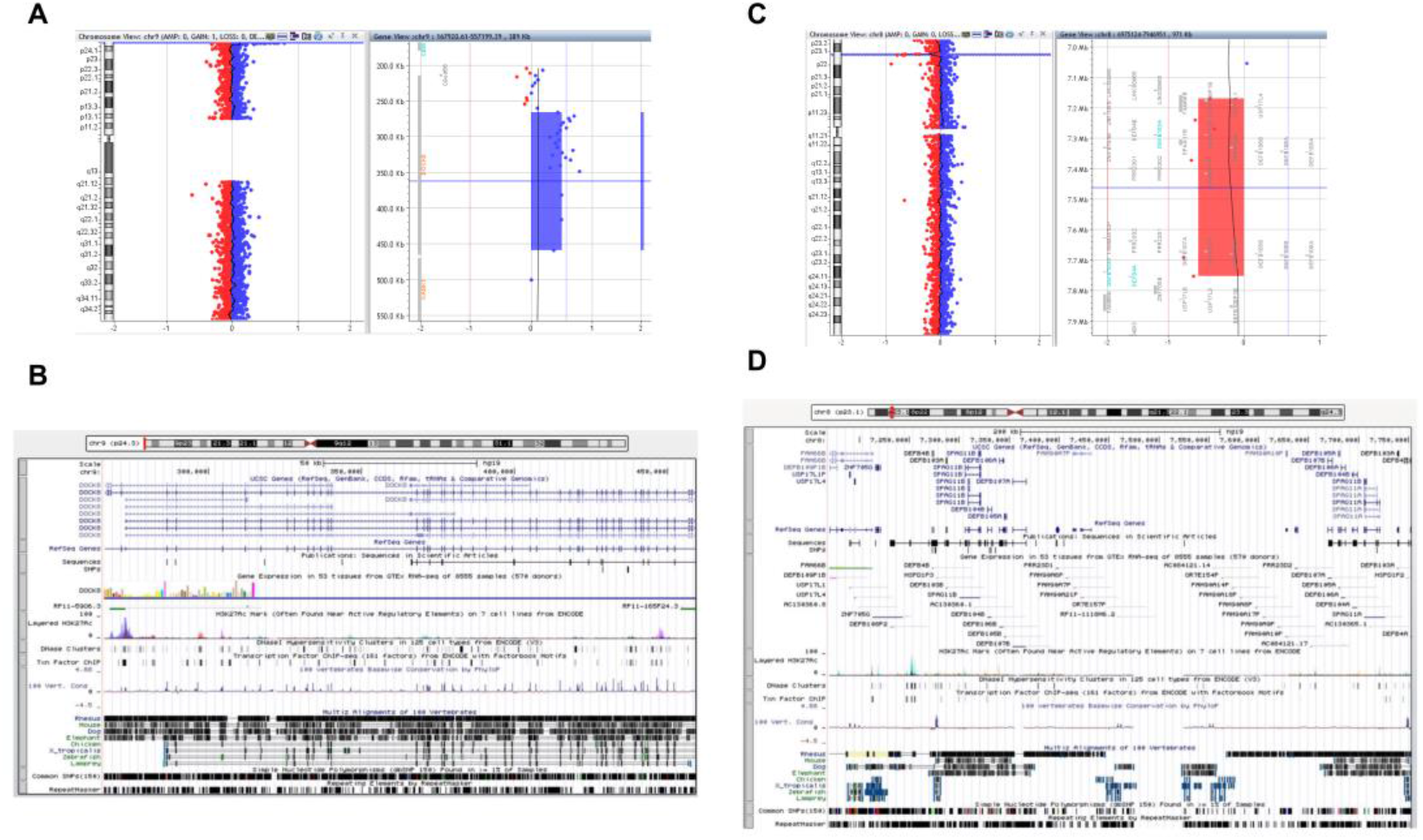
Chromosomal alterations found in our proband. A. Screen capture of the array-CGH of the proband’s chromosome 9 showing the microduplication at 9p24.3. B. Screen capture of the UCSC Genome Browser (https://genome.ucsc.edu/) showing the gene partially duplicated in the proband. C. Screen capture of the array-CGH of the proband’s chromosome 8 showing the microdeletion at 8p23.1. D. Screen capture of the UCSC Genome Browser (https://genome.ucsc.edu/) showing the genes partially deleted in the proband.

In order to delve into the molecular causes of the speech and language deficits exhibited by the child, we conducted several silico analyses. First, we surveyed the literature looking for patients with language problems in which any of the genes deleted or duplicated in our proband is affected. None of the genes located within the deleted region in chromosome 8 has been associated to language deficits. Defensins are peptides with a broad-spectrum antimicrobial activity and with no known role in brain development or function, beyond host defense against central nervous system pathogenesis (Hao et al., 2001). Defensin CNVs are well-known, but they have been related mostly to differential susceptibility to infection and disease (Linzmeier and Ganz, 2005). Likewise, the two sperm associated antigen proteins (SPAG11A and SPAG11B) encode β-defensins involved in sperm maturation (Chan and Yang, 2005; Hu et al., 2014), with no known role in brain development either. The two proline rich domain proteins (PRR23D1 and PRR23D2), the two ubiquitin peptidase-like proteins (USP17L1P and USP17L4), and the zinc finger protein (ZNF705G) play unknown roles. On the contrary, our proband exhibits several of the neurological, cognitive, and behavioral features of *DOCK8*-deficient patients, as well as individuals with duplications of the 9p24.3 region encompassing *DOCK8* (Table 3).

**Table 3.** Summary table with the most relevant clinical features of our proband compared to patients with *DOCK8* deficiency (according to Griggs et al., 2008 and Biggs et al. 2017, and Engelhardt et al., 2017) and with duplications affecting or encompassing *DOCK8* (up to 1Mb) (according to DECIPHER). The observed features are marked with x.

| **Clinical features of *DOCK8* deficiency** |  |
| --- | --- |
| Cognitive/behavioural features |  |
| Mental retardation | x |
| Absence of speech/language impairment | x |
| Physical features |  |
| Dysmorphic facial features | x |
| Increased nose width |  |
| Retained primary teeth |  |
| High palate |  |
| Other medical problems |  |
| Neurological complications |  |
| Skin disease (eczema, abscesses) |  |
| Lung disease/abnormalities (pneumonia, bronchiectasis) |  |
| Atopy (asthma, allergies) |  |
| Autoimmune disease |  |
| Malignancies |  |
| Hyperflexibility |  |
| High susceptibility to superficial, localized, and invasive infections |  |

**Table 3.** Summary table with the most relevant clinical features of our proband compared to patients with *DOCK8* deficiency (according to Griggs et al., 2008 and Biggs et al.
| **Clinical features of *DOCK8* duplications (up to 1Mb)** |  |
| --- | --- |
| Cognitive/behavioural features |  |
| Intellectual disability | x |
| Global developmental delay | x |
| Autism |  |
| Behavioral abnormality | x |
| Delayed speech and language development | x |
| Neurological findings |  |
| Motor delay | x |
| Epileptic spasms |  |
| Febrile seizures |  |
| Physical features |  |
| Macrocephaly |  |
| Dolichocephaly |  |
| Dysmorphic facial features (long philtrum) | x |

Additionally, we surveyed DECIPHER (https://decipher.sanger.ac.uk/) to find other patients with deletions and duplications that are similar or smaller than the fragments deleted or duplicated in our proband, that might help associate his language problems to specific genes. We found that microdeletions in 8p23.1 are usually reported as benign (patients 298702, 285986, 298151), of uncertain pathogenicity (patient 323189), or of unknown pathogenicity (patients 317632 and 317616). DECIPHER patients exhibiting developmental delay (298702, 285986, and 298151) bear other CNVs in other chromosomes that are reported as pathogenic and that seemingly account for their problems in the cognitive/behavioral domains. Three DECIPHER patients (323189, 317632, and 317616) bear unique deletions in chromosome 8 that are similar to the one found in our patient, but neither of them is reported to present with language delay, speech problems, cognitive disabilities, mental retardation, or learning impairment. By contrast, DECIPHER patients with duplications in 9p24.3 encompassing *DOCK8* frequently present with intellectual disability, developmental delay, behavioral abnormality, and/or delayed speech and language development, even in the absence of additional CNVs (e.g. patients 328340 and 251154) (Figure 3).

**Figure 3.**
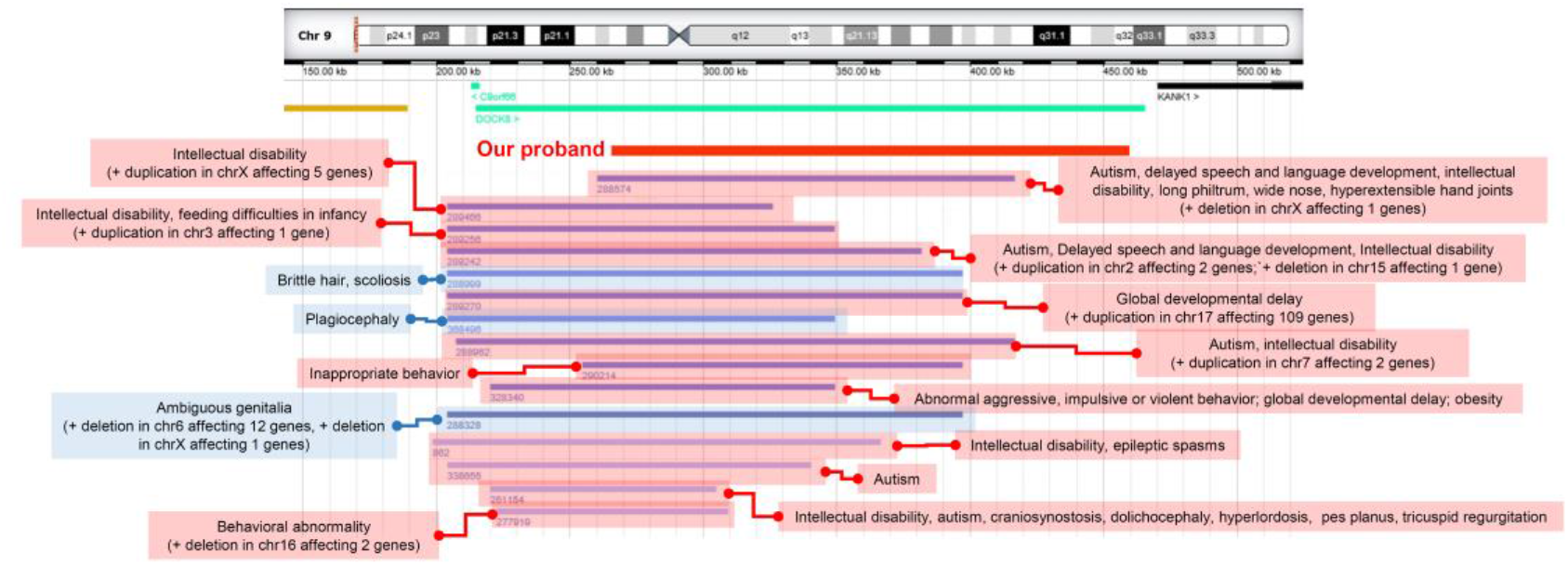
DECIPHER patients of interest bearing chromosomal duplications at 9p24.3 with similar sizes than the one found in the proband. Patients with cognitive deficits are highlighted in red, cases with no reported cognitive problems are highlighted in blue.

Finally, we used String 10.5 (www.string-db.org) for uncovering potential functional links between the genes deleted and duplicated in our proband and genes important for language. We focused on candidate genes for i) prevalent language disorders (developmental dyslexia (DD) and specific language impairment (SLI), as listed by Paracchini et al., 2016, Pettigrew et al 2016, and Chen et al. 2017); and ii) language evolution, as discussed by Boeckx and Benítez-Burraco (2014a,b) and Benítez-Burraco and Boeckx (2015) (many of the genes belonging to this second group are also candidates for language dysfunction in broader cognitive disorders, particularly, ASD and SZ, as discussed in detail in Benítez-Burraco and Murphy (2016), Murphy and Benítez-Burraco (2016) and Murphy and Benítez-Burraco (2017)). The whole set of genes considered in this analysis is listed in the Supplemental file 3. String 10.5 predicts physical and functional associations between proteins relying on different sources (genomic context, high-throughput experiments, conserved coexpression, and text mining) (Szklarczyk et al., 2015). Because we were interested in finding robust functional links, we restricted our search to significant protein interactions gathered from protein-protein interaction databases and curated databases (Biocarta, BioCyc, GO, KEGG, and Reactome). String did not predict any functional link between any of the proteins coded by the genes within the 8p23.1 deleted region and the proteins coded by candidates for language disorders and/or evolution (data not shown). By contrast, it predicted a robust direct link between DOCK8 and CDC42, as well as robust indirect links with nearly 30 other candidates for language disorders and/or evolution, including SLIT1, SLIT2, and ROBO1, which play a key role in the externalization of language (speech) (see Boeckx and Benítez-Burraco, 2014b for review) (Figure 4).

**Figure 4.**
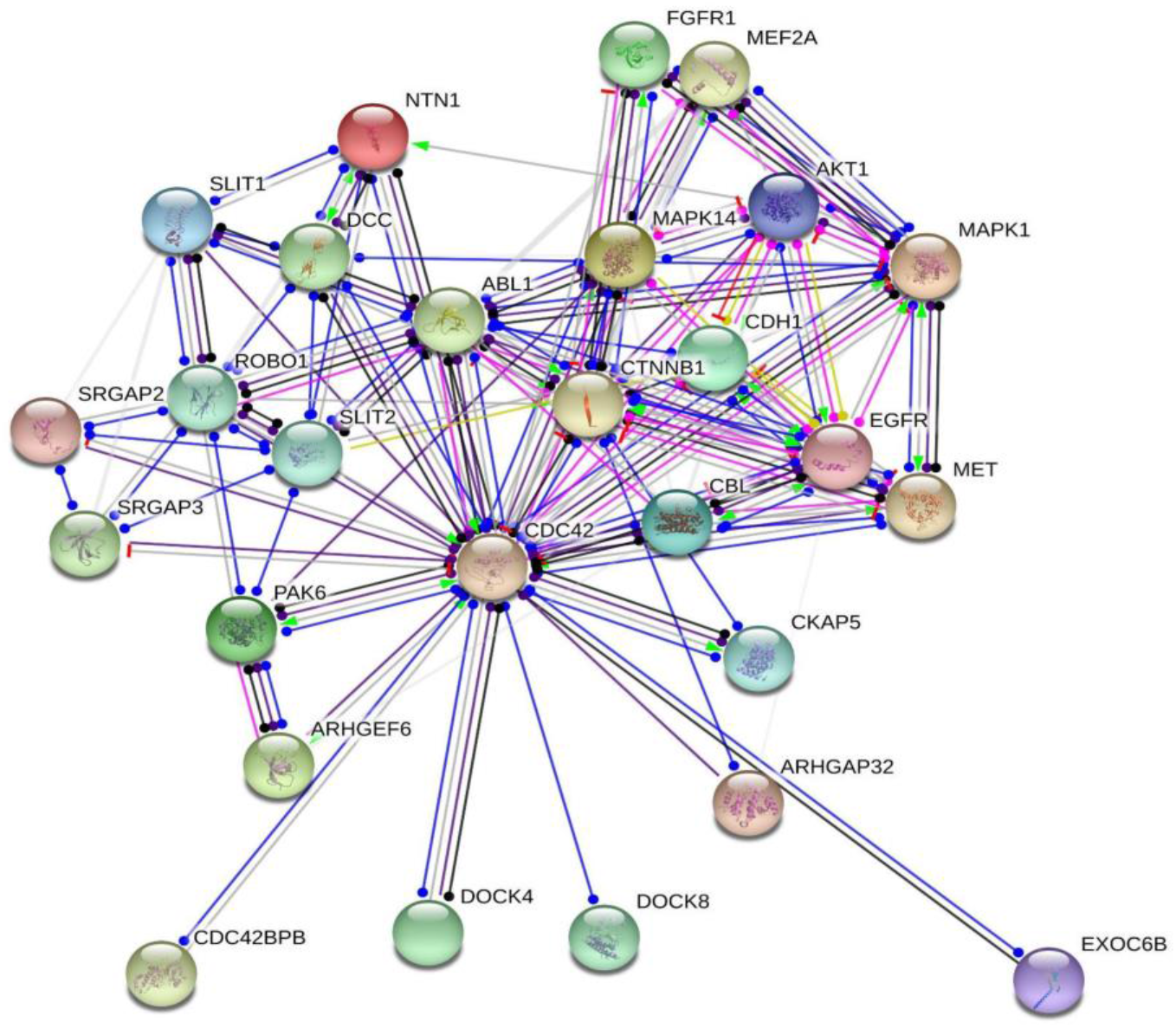
Interaction network of the DOCK8 protein with proteins encoded by core candidates for language development and evolution. The network was drawn with String (version 10.5; Szklarczyk et al. 2015) license-free software (http://string-db.org/), using the molecular action visualization. It includes DOCK8 and the products of nearly 30 strong candidates for language development/evolution. Colored nodes symbolize gene/proteins included in the query (small nodes are for proteins with unknown 3D structure, while large nodes are for those with known structures). The color of the edges represents different kind of known protein-protein associations. Green: activation, red: inhibition, dark blue: binding, light blue: phenotype, dark purple: catalysis, light purple: post-translational modification, black: reaction, yellow: transcriptional regulation. Edges ending in an arrow symbolize positive effects, edges ending in a bar symbolize negative effects, whereas edges ending in a circle symbolize unspecified effects. Grey edges symbolize predicted links based on literature search (co-mentioned in PubMed abstracts). Stronger associations between proteins are represented by thicker lines. The medium confidence value was .0400 (a 40% probability that a predicted link exists between two enzymes in the same metabolic map in the KEGG database: http://www.genome.jp/kegg/pathway.html). The diagram only represents the potential connectivity between the involved proteins, which has to be mapped onto particular biochemical networks, signaling pathways, cellular properties, aspects of neuronal function, or cell-types of interest.

Besides in silico analyses, we also conducted in vitro analyses. Specifically, we performed microarray analyses of blood samples from the proband to determine whether he exhibited altered patterns of gene expression that may account for the observed symptoms. We used his healthy parents as controls. The results of the microarrays are shown in the Supplemental file 4. First, we checked whether the genes within the deleted or duplicated chromosomal fragments were differentially expressed in the proband compared to his unaffected parents (Figure 5A). We found that *DOCK8* was slightly upregulated in the proband. Regarding the genes within the deleted fragment in 8p23.1, we found that several genes encoding defensins were downregulated, particularly, *DEFB105A, DEFB105B, DEFB106B, DEFB107A* and *DEFB107B*. By contrast, *SPAG11B* and *ZNF705G* were slightly upregulated.

**Figure 5.**
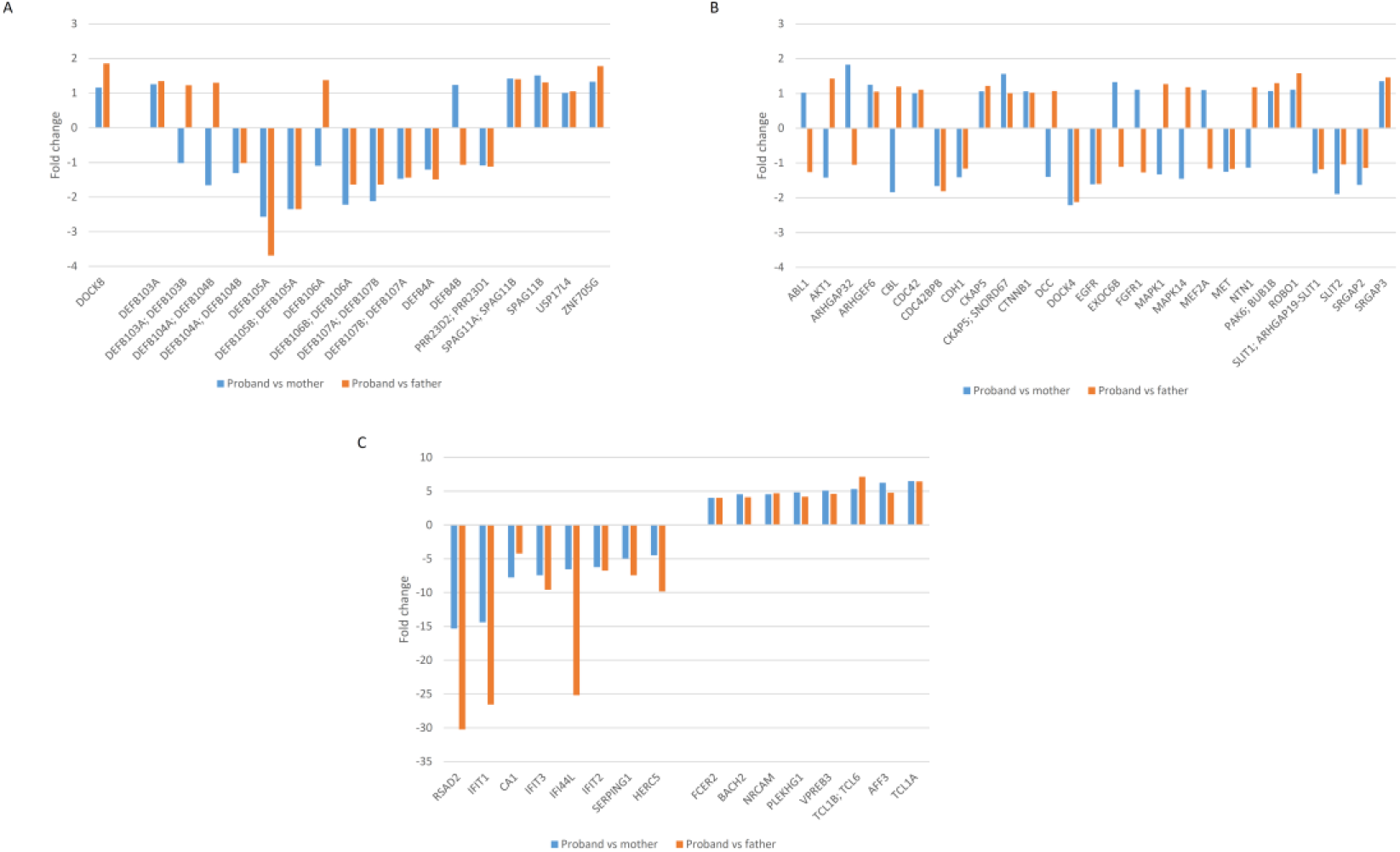
Variation in the expression levels of genes of interest in the proband’s blood compared to her healthy parents. A. Fold changes in the expression levels of *DOCK8*, the only gene within the region duplicated at 9p24.3 in the proband (left), and of the genes within the region deleted at 8p23.1 (right). B. Fold changes in the expression levels of predicted functional partners of DOCK8 with a known role in language development, language impairment, and/or language evolution. C. Genes exhibiting fold changes > 4 in the proband compared to both healthy parents.

Second, we checked whether the genes encoding proteins predicted by String 10.5 to interact with DOCK8 were differentially expressed in the boy compared to his healthy parents (Figure 5B). For most genes we found no significant differences. However, some of them were downregulated (*CDC42BPB, DOCK4, EGFR*), whereas *SRGAP3* was upregulated in the proband. Minor differences were found in *ROBO1* (slightly upregulated) and SLIT factors (slightly downregulated), as well as in *FOXP2* (slightly upregulated; see Supplemental file 4).

Finally, we searched for additional candidates for the proband’s language deficits, looking for genes exhibiting strong fold changes (FC) in our subject (i.e. with FC > 4 compared to both unaffected parents). We found several strongly downregulated genes (*HERC5, SERPING1, IFIT2, IFI44L, IFIT3, CA1, IFIT1* and *RSAD2*), as well as several strongly upregulated genes (*FCER2, BACH2, NRCAM, PLEKHG1, VPREB3, TCL1B, TCL6*, *AFF3* and *TCL1A*) (Figure 5C).

deleted fragment in 8p23.1, neither of them are expressed in the brain at any significant level (Figure 6A). By contrast, most of the potential interactors of DOCK8 with a known play in language development and/or evolution are highly expressed in the brain (with the exception of *CDH1, DCC*, and *MET*). In fact, they are expressed in the brain at higher levels than in the blood (with the exception of *MAPK14*). Specifically, most of them are preferentially expressed in the cerebellum and to some extent in the cortex and the basal ganglia (Figure 6B). Finally, with regards to the genes that are strongly downregulated in our proband, we found that many of them were highly expressed in most brain areas, particularly, *IFT1, IFT2, IFT3*, and *SERPING1*. Regarding the genes found strongly upregulated in the boy, *NRCAM* is highly expressed in all brain areas, whereas *AFF3* and *PLEKHG1* are preferentially expressed in the cerebellum and the spinal cord, respectively.

**Figure 6.**
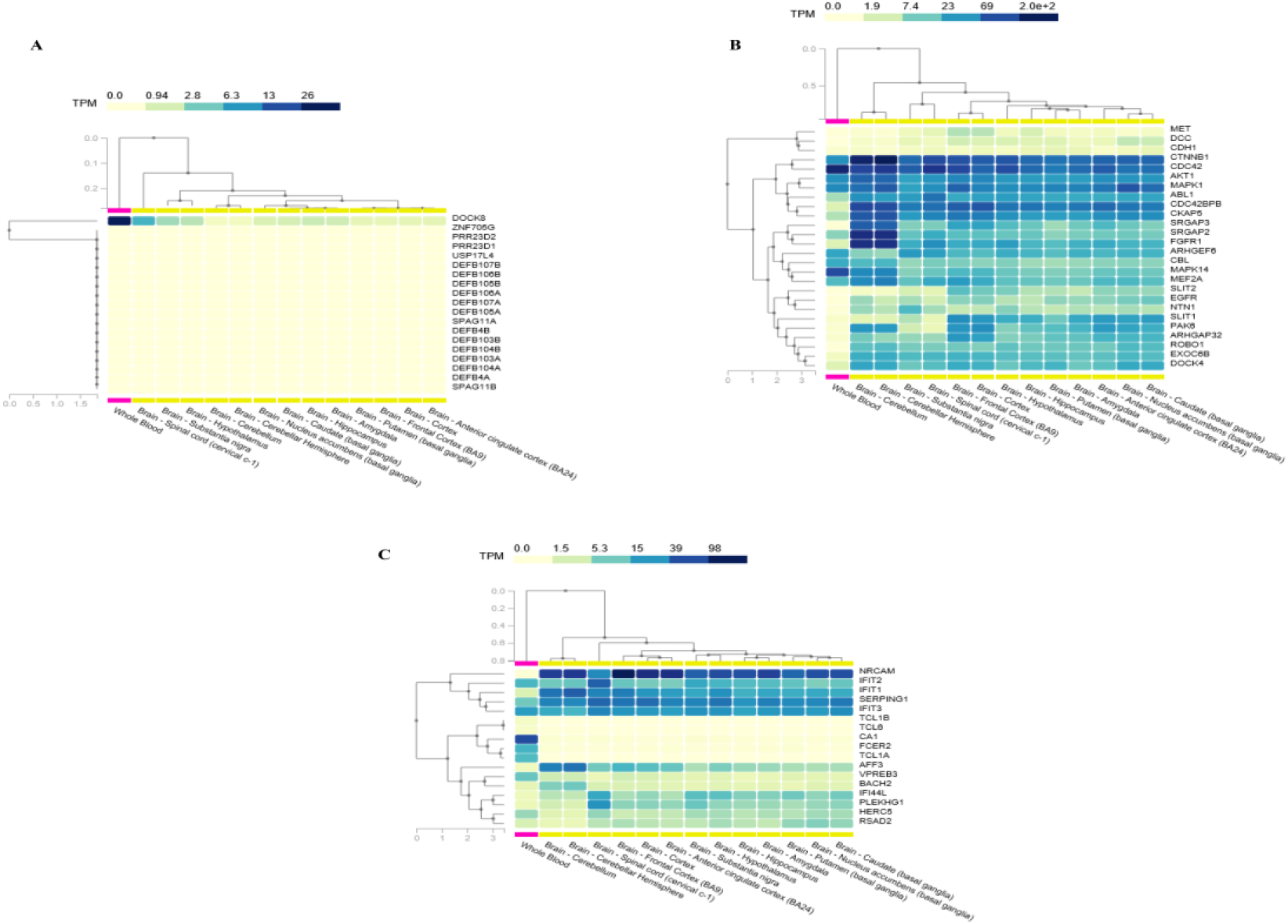
Expression pattern in the blood and the brain of genes of interest. A. Expression profiles of DOCK8 and of the genes within the region deleted at 8p23.1 in the proband. B. Expression profiles of predicted functional partners of DOCK8 with a known role in language development, language impairment, and/or language evolution. C. Expression profiles of genes exhibiting fold changes > 4 in the proband compared to both healthy parents. Expression levels of the genes were retrieved from the Genotype-Tissue Expression (GTEx) project (GTEx Consortium, 2013) (https://www.gtexportal.org/home/). Statistical analysis and data interpretation were performed by The GTEx Consortium Analysis Working Group using the. Legend: TPM, transcripts per million.

## DISCUSSION

The generalized application of next generation sequencing facilities, particularly array-CGH, to the genetic examination of subjects with cognitive and language disorders has resulted in a growing list of genes and chromosomal regions associated to these conditions. Nonetheless, in most cases, no robust genotype-to-phenotype links have been established and no molecular causative factors have been identified to date. In this paper we have provided a detailed description of the cognitive and linguistic features of a boy with a microdeletion in 8p23.1 and a microduplication in 9p24.3. It is our contention, however, that his abnormal phenotype, and particularly, his problems with language, result from the partial duplication of the *DOCK8* gene in 9p24.3. On the one hand, the boy exhibits many of the physical, neurological, cognitive, behavioral, and language features of people with CNVs encompassing this gene (Table 3 and Figure 3). Moreover, we have found no evidence in the literature or in patient databases of the involvement of any of the genes within the deleted region in 8p23.1. in language development, or in cognitive and behavioral aspects of relevance for language function. Finally, as we discuss below in more detail, *DOCK8* is functionally connected with many genes that are robust candidates for language development, impairment, and/or evolution. Although alterations of *DOCK8* are known to impact on language abilities and although the gene is known to play a role in brain development and function, we still lack a detailed characterization of the language phenotype resulting from *DOCK8* alterations. One important reason is that the linguistic evaluation of patients usually involves psychological tests only. In our work, we have conducted a fine-grained analysis of the language strengths and deficits of our subject using a battery of specific diagnostic tools, but also examining samples of naturalistic speech, looking for structural and functional anomalies.

In the expressive domain, our proband shows signals of a slightly delayed phonological development, characterized by sound substitutions, simplification of consonant clusters, assimilations, omissions, weak syllable deletions, and problems with syllable segmentation. His score in the Phonological Awareness subtest of the CELF-4 reinforces this view. Additionally, his ability to assemble morphemes into words and words into sentences is also impaired. His MLUw is shorter than expected according to his chronological age. Some functional components of language, like inflectional affixes or prepositions, are particularly problematic for him. He scored very low in tasks assessing his ability to compose sentences and to retrieve and process verbal information. These difficulties with utterance production, as well as the reduced intelligibility resulting from his articulatory and phonological problems, explain his frequent repetitions of words and phrases, and ultimately, his perseverative discourse. Receptive language is also affected, as evidenced by the child’s low scores in tests evaluating his receptive vocabulary and his comprehension of the grammar, as well as his echolalic responses during conversation. Complex sentences such as those with non-canonical word order (OVS), passives, and with relative clauses, are particularly problematic for him. Finally, he exhibits a good semantic knowledge at the word level, but has problems with aspects of sentence semantics involving structural aspects.

In our opinion, the observed deficits mostly result from the impairment of more basic cognitive abilities, specifically, working memory. On the one hand, the child’s nonverbal IQ is suggestive of a moderate mental retardation. On the other hand, he scored very low in tasks evaluating working memory (e.g. the WMI of the CELF-4) and exhibited difficulties in sustaining focus. Deficits in working memory have been reported to result in learning disabilities, poor phonological awareness, and impaired sequential processing in subjects with other chromosomopathies such as Fragile X syndrome (Johnson-Glenberg, 2008). In particular, problems with sequential processing can contribute to explain the syntactic perseveration observed in our proband. At the same time, we cannot rule out the possibility that his problems with language are exacerbated because of his reduced executive function and/or attention, as the child presents with lack of inhibition and behaves impulsively and inappropriately to age. The co-occurrence of deficits in attention, working memory, and inhibition together with language problems has been reported in other clinical conditions, like Fragile X syndrome (Artigas-Pallarés et al., 2001). That said, in spite of these pervasive problems with structural aspects of language, the child has mastered some key aspects of language use, like the dynamics of turn-taking and compensatory behaviors aimed to circumvent expressive difficulties. His willingness to communicate is remarkable. Still, he frequently violates conversational principles (like the maxims of quality and relationship) and seems unable to understand the logics of conversation. These circumstances make his discourse incomplete, awkward, and incoherent at times. Because his semantic knowledge is spared, we contend that these problems with language use might be caused by a deviant reasoning, lack of attention, and reduced encyclopedic knowledge (i.e. awareness of the common ground shared with his interlocutors). Limited memory abilities, ultimately boiling down to his working memory deficit, seemingly contribute to these problems.

To sum up, our proband exhibits selected deficits in several structural components of language (notably, inflectional morphology, complex syntax, and sentence semantics), both in the expressive and receptive domains, together with a pragmatic impairment, which seemingly result from his intellectual disability, and particularly, from his severe deficit in working memory.

Regarding the molecular causes of the observed symptoms, we contend that they might arise from an abnormal interaction in the brain between DOCK8 and CDC42 resulting in the dysregulation of several key genes involved in language development and/or evolution. We cannot rule out the possibility that changes in the expression levels of selected genes outside the duplicated region and not interacting with DOCK8 also contribute to the problems exhibited by the proband.

In the blood, DOCK8 activates CDC42 at the leading edge membrane of migrating cells, with this activation promoting changes in cell morphology that are needed for proper 3-D cell migration (Harada et al., 2012). CDC42 regulates neural crest cell proliferation (Fuchs et al., 2009) and cortical interneuron migration (Katayama et al. 2013), and CDC42 signaling is involved as well in dendritic spine growth and function (Datta et al., 2015). *CDC42* polymorphisms have been associated to increased SZ risk (Gilks et al., 2012). As shown in Figure 4, CDC42 interacts with nearly 30 candidates for language development, evolution, and/or impairment. Most of these genes do not show fold changes in our proband compared to his healthy parents, but some of them appear slightly down- or upregulated. In view of the speech and language problems exhibited by the child, *SLIT2* (slightly downregulated) and *ROBO1* (slightly upregulated) are of particular interest. Slit2 reduces Cdc42 activity in living growth cones (Myers et al. 2012). In turn, depletion of Robo1 prevents Slit2 inhibition of Cdc42 activity (Yiin et al., 2009). The SLIT/ROBO pathway has been claimed to play an important role in the externalization of language (i.e. speech) (see Boeckx and Benítez-Burraco 2014b for details). Specifically, the SLIT-mediated interaction between FOXP2/foxp2 (also found slightly upregulated in our proband and reviewed above) and ROBO1/robo1 is critical to the establishment of vocal learning pathways across species (Pfenning et al., 2014).

Another gene of interest is *CDC42BPB*, which is found downregulated in our proband. This gene encodes an effector of CDC42 and is regulated by FOXP2 (Spiteri et al., 2007). *FOXP2* is a strong candidate for speech and language impairment (Vargha-Khadem, et al., 2005; Kurt et al., 2012; Graham et al., 2015), playing a key role in neurogenesis, neuron differentiation, and neuron migration in brain areas involved in language processing (Tsui et al., 2013; Chiu et al., 2014; García-Calero et al., 2016). Pathogenic mutations in humans have been proven to affect auditory-motor association learning when mimicked in mice (Kurt et al., 2012). Specifically, the gene seems to facilitate the transition from declarative (i.e. less automatized) to procedural (i.e. more automatized) performance (Schreiweis et al., 2014; Chandrasekaran et al., 2015).

Interestingly too, *SRGAP3* is found upregulated in our proband. This gene has been associated to absence of speech co-occurring with severe mental retardation (Endris et al., 2002), as well as to schizophrenia (Wilson et al., 2011; Waltereit et al., 2012). *SRGAP3* is involved in cytoskeletal reorganization and contributes to different neurodevelopmental processes, including neuronal differentiation and neurite outgrowth, which may be required for normal cognitive function (Endris et al., 2011; Bacon et al., 2013; Ma et al., 2013). SRGAP3 interacts with ROBO1, thus contributing to modulate the SLIT/ROBO pathway (Wong et al., 2001).

To finish, it is also of interest the finding that *DOCK4* and *EGFR* are downregulated in our proband. A microdeletion of *DOCK4* has been suggested to contribute to dyslexia, as well as to ASD (Pagnamenta et al., 2010). Concerning *EGFR*, this gene interacts with *VCAN*, a gene involved in neuronal attachment, neurite outgrowth, and synaptic transmission (Xiang et al., 2006).

Lastly, it is worth considering as well the biological roles played by the genes that are more strongly dysregulated in the blood of our proband compared to his healthy parents. Among the genes found strongly downregulated, most of them are involved in antiviral response (*RSAD2, IFIT1, IFIT2, IFIT3, IFI44L). CA1* plays diverse biological roles, but although it is highly expressed in the blood, it is not expressed in the brain (Figure 6). Accordingly, the only potential candidate for the cognitive and language deficits exhibited by our proband is *SERPING1*. This gene encodes a plasma protein involved in the regulation of the complement cascade. This cascade has been related to synaptic plasticity, as the knockdown of *Serping1* in mice impairs synapse formation (Ren et al., 2020). The gene has been also linked to cortical development, particularly, to neuronal migration (Gorelik et al., 2017).

The genes found strongly upregulated in the proband are of more interest in view of his language and cognitive phenotype. Several of them (*FCER2, TCL1A, TCL1B, TCL6*) are not expressed in the brain or are expressed at very low levels, and have been associated to conditions not involving cognitive deficits, like cancer (*TCL1A*, *TCL1B*, *TCL6*) or asthma (*FCER2*). Interestingly though, *FCER2* and *TCL1B* have been found to be dysregulated in the blood of patients with Parkinson’s disease (Infante et al., 2015). Regarding the genes that are expressed in the brain, *NRCAM* is the most interesting candidate. This gene is highly expressed in most brain areas (particularly, the cortex and the thalamus) and encodes a neuronal cell adhesion protein with attested roles in neuron-neuron adhesion, as well as axonal cone growth and axonal guidance, being ultimately involved in the organization of neural circuits during brain development (Sakurai, 2012). Variants of this gene have been associated with ASD, particularly, with the subtype co-occurring with obsessive-compulsive behavior (Sakurai et al., 2006), as well as with addictive behavior because of its regulatory role in glutamatergic and GABAergic innervation (Ishiguro et al., 2014; Ishiguro et al., 2019). All this evidence suggests that *NRCAM* might play some important role in the reward circuitry, in part through its involvement in the fasciculation of axon fibers in the amygdala (Mohan et al., 2019). In view of the attention problems exhibited by our proband, it is interesting that *Nrcam-null* mice show decreased sociability and learning deficits (Moy et al., 2009), as well as altered impulsivity (Matzel et al., 2008). Two other genes of interest, that are also strongly upregulated in our proband and also expressed in the brain, are *AFF3* and *BACH*2. They are mostly expressed in the cerebellum, particularly, after birth (Supplemental file 5). *AFF3* encodes a nuclear transcription factor required for normal cellular migration in the developing cortex (Moore et al., 2014). Silencing of this gene results in intellectual disability (Metsu et al., 2014). Regarding *BACH2*, it encodes a transcription factor which is part of a regulatory network involved in the development of cortico-striatal projections within the telencephalon, also encompassing *FOXO1* and *ISL1* (Waclaw et al., 2017). *FOXO1* is a target of FOXP2 (Vernes et al., 2011), whereas the FOXO1 protein is phosphorylated by DYRK1A (Huang and Tindall, 2007), associated to mental retardation and absence of speech (Van Bon et al., 2011; Courcet et al., 2012) and involved in learning and memory (Hämmerle et al. 2003). In turn, FOXO1 upregulates *RELN* (Daly et al. 2004), a candidate for ASD and for lissencephaly with language loss (Hong et al., 2000; Wang et al., 2014). Finally, *PLEKHG1* encodes an activator of CDC42 (Reinhard et al., 2019) and has been associated to panic disorder (Otowa et al., 2009) and to white matter hyperintensities (Traylor et al., 2019).

## CONCLUSIONS

Although the exact molecular causes of the language problems (and other cognitive, behavioral and even motor deficits) observed in our proband remain to be fully elucidated, we hypothesize that his distinctive features might result from the altered expression of selected genes involved in procedural learning, particularly, some components of the SLIT/ROBO/FOXP2 network, which are strongly related to the development and evolution of language. Dysregulation of specific components of this network can be hypothesized to result in turn from an altered interaction between DOCK8, affected by the microduplication in 9p24.3 borne by our proband, and CDC42, acting as the hub component of the network encompassing language-related genes. Still, some genes found strongly upregulated in the subject, and not related to these genes, can contribute to the observed problems in the language domain, as well as to specific features of the proband, particularly, impulsivity.

Although this hypothesis needs to be properly tested, we hope that our findings contribute to a better understanding of the language, cognitive, and behavioral phenotype resulting from CNVs of the distal region of the short arm of chromosome 9. A more detailed knowledge of the neurobiological basis of the deficits (and strengths) exhibited by patients is necessary for improving (psycho)pedagogical strategies aimed to ameliorate their problems. This is particularly true of low-prevalent clinical conditions, such as this kind of CNVs, which are mostly understudied and for which optimized intervention tools are not usually available.

## ACKNOWLEDGMENTS

We would like to thank the proband and his family for their participation in this research. We wish to thank also Dr. Montserrat Barcos Martínez and Dr. Isabel Espejo Portero, from the “Reina Sofía” hospital in Córdoba, for giving us access to the patient and his clinical history. Preparation of this work was supported by funds from the Spanish Ministry of Economy and Competitiveness (grant number FFI2016-78034-C2-2-P [AEI/FEDER,UE] to Antonio Benítez-Burraco, with MFU and SJR as members of the project).

## ETHICS

Ethics approval for this research was granted by the Comité Ético del Hospital “Reina Sofía”. Written informed consent was obtained from the proband’s parents for conducting the psycholinguistic evaluations and the molecular analyses, and for publicizing this case report and all the accompanying tables and images in scientific journals and meetings.

## AUTHORS’ CONTRIBUTION

ABB conceived the paper, performed the molecular experiments, analyzed the genomic/genetic data, and wrote the paper. MFU and SJR conducted the clinical evaluation of the proband, analyzed the psycholinguistic data, and wrote the paper. The three authors approved the final version of the manuscript.

## SUPPLEMENTAL DATA

**Supplemental file 1.** Tables with the scores obtained by the proband in the standardized tests used for her assessment.

**Supplemental file 2.** Transcriptions of the conversational exchanges in natural settings used for the analyses of the proband’s language in use.

**Supplemental file 3.** The whole set of genes important for language used for the in silico analysis with String 10.5.

**Supplemental file 4.** Data resulting from the microarray analyses comparing gene expression levels in the blood of the proband and his healthy parents.

**Supplemental file 5.** Developmental expression profiles of genes of interest. These genes include *DOCK8*, the only gene within the region duplicated at 9p24.3 in the proband; the genes within the region deleted at 8p23.1; predicted functional partners of DOCK8 with a known role in language development, language impairment, and/or language evolution; and genes exhibiting fold changes > 4 in the proband compared to both healthy parents, as discussed in the paper. The expression data are from the Human Brain Transcriptome Database (http://hbatlas.org/). Six different brain regions are considered: the cerebellar cortex (CBC), the mediodorsal nucleus of the thalamus (MD), the striatum (STR), the amygdala (AMY), the hippocampus (HIP) and 11 areas of neocortex (NCX).

## REFERENCES

Allen-Brady K, Miller J, Matsunami N, Stevens J, Block H, Farley M, Krasny L, Pingree C, Lainhart J, Leppert M, McMahon WM, Coon H (2009) A high-density SNP genome-wide linkage scan in a large autism extended pedigree. Mol Psychiatry 14: 590–600

Artigas-Pallarés J, Brun C, Gabau E (2001). [Medical and neuropsychological aspects of fragile X syndrome]. Revista de Neurología Clínica 2(1): 42–54.

Bacon C, Endris V, Rappold GA (2013) The cellular function of srGAP3 and its role in neuronal morphogenesis. Mech Dev. 130: 391–5

Benítez-Burraco A, Boeckx C. (2015) Possible functional links among brain- and skull-related genes selected in modern humans. Front Psychol. 6: 794.

Benítez-Burraco A, Murphy E. (2016) The oscillopathic nature of language deficits in autism: from genes to language evolution. Front Hum Neurosci 10: 120.

Biggs CM, Keles S, Chatila TA (2017) DOCK8 deficiency: Insights into pathophysiology, clinical features and management. Clin Immunol 181: 75–82.

Bishop DVM (2003) Test for reception of grammar: TROG-2. London: Harcourt Assessment

Boeckx C, Benítez-Burraco A (2014a) The shape of the human language-ready brain. Front Psychol 5: 282.

Boeckx C, Benítez-Burraco A. (2014b) Globularity and language-readiness: Generating new predictions by expanding the set of genes of interest. Front Psychol. 5: 1324.

Broides A, Mandola AB, Levy J, Yerushalmi B, Pinsk V, Eldan M, Shubinsky G, Hadad N, Levy R, Nahum A, Ben-Harosh M, Lev A, Simon A, Somech R (2017) The clinical and laboratory spectrum of dedicator of cytokinesis 8 immunodeficiency syndrome in patients with a unique mutation. Immunol Res. 65: 651–657

Cattell RB, Cattell AKS (1994) Tests de Factor «g», Escalas 2 y 3. Madrid: TEA Ediciones, S. A.

Chan HC, Zhang YL (2005) Epididymial defensins and sperm maturation. Andrologia 37: 200–1.

Chandrasekaran B, Yi HG, Blanco NJ, McGeary JE, Maddox WT (2015) Enhanced procedural learning of speech sound categories in a genetic variant of FOXP2. Neurosci. 35(20):7808–12. doi: 10.1523/JNEUROSCI.4706-14.2015.

Chen XS, Reader RH, Hoischen A, Veltman JA, Simpson NH, Francks C, Newbury DF, Fisher SE (2017) Next-generation DNA sequencing identifies novel gene variants and pathways involved in specific language impairment. Sci Rep. 7: 46105.

Chiu Y-C, Li MY, Liu YH, Ding JY, Yu JY, Wang TW (2014) Foxp2 regulates neuronal differentiation and neuronal subtype specification. Dev Neurobiol. 74: 723–738.

Conners CK (2008) Conners Comprehensive Behavior Rating Scales (Conners CBRS). Toronto: MHS

Conti-Ramsden G (1996) CLAN (Computerized Language Analysis). Child Lang Teach Ther. 12: 345–349.

Coon H, Villalobos ME, Robison RJ, Camp NJ, Cannon DS, Allen-Brady K, Miller JS, McMahon WM (2010) Genome-wide linkage using the Social Responsiveness Scale in Utah autism pedigrees. Mol Autism. 1: 8.

Courcet JB, Faivre L, Malzac P, Masurel-Paulet A, Lopez E, Callier P, Lambert L, Lemesle M, Thevenon J, Gigot N, Duplomb L, Ragon C, Marle N, Mosca-Boidron AL, Huet F, Philippe C, Moncla A, Thauvin-Robinet C (2012) The DYRK1A gene is a cause of syndromic intellectual disability with severe microcephaly and epilepsy. J Med Genet. 49(12): 731–6. doi: 10.1136/jmedgenet-2012-101251.

Daly C, Wong V, Burova E, Wei Y, Zabski S, Griffiths J, Lai KM, Lin HC, Ioffe E, Yancopoulos GD, Rudge JS (2004) Angiopoietin-1 modulates endothelial cell function and gene expression via the transcription factor FKHR (FOXO1). Genes Dev. 18(9):1060–71. doi: 10.1101/gad.1189704.

Datta D, Arion D, Corradi JP, Lewis DA (2015) Altered expression of CDC42 signaling pathway components in cortical layer 3 pyramidal cells in schizophrenia. Biol Psychiatry 78: 775–85.

De Rubeis S, He X, Goldberg AP, Poultney CS, Samocha K, Cicek AE, Kou Y, Liu L, Fromer M, Walker S, Singh T, Klei L, Kosmicki J, Shih-Chen F, Aleksic B, Biscaldi M, Bolton PF, Brownfeld JM, Cai J, Campbell NG, Carracedo A, Chahrour MH, Chiocchetti AG, Coon H, Crawford EL, Curran SR, Dawson G, Duketis E, Fernandez BA, Gallagher L, Geller E, Guter SJ, Hill RS, Ionita-Laza J, Jimenz Gonzalez P, Kilpinen H, Klauck SM, Kolevzon A, Lee I, Lei I, Lei J, Lehtimäki T, Lin CF, Ma’ayan A, Marshall CR, McInnes AL, Neale B, Owen MJ, Ozaki N, Parellada M, Parr JR, Purcell S, Puura K, Rajagopalan D, Rehnström K, Reichenberg A, Sabo A, Sachse M, Sanders SJ, Schafer C, Schulte-Rüther M, Skuse D, Stevens C, Szatmari P, Tammimies K, Valladares O, Voran A, Li-San W, Weiss LA, Willsey AJ, Yu TW, Yuen RK; DDD Study; Homozygosity Mapping Collaborative for Autism; UK10K Consortium, Cook EH, Freitag CM, Gill M, Hultman CM, Lehner T, Palotie A, Schellenberg GD, Sklar P, State MW, Sutcliffe JS, Walsh CA, Scherer SW, Zwick ME, Barett JC, Cutler DJ, Roeder K, Devlin B, Daly MJ, Buxbaum JD (2014) Synaptic, transcriptional and chromatin genes disrupted in autism. Nature 515: 209–15.

Diez-Itza E, Martínez V, Pérez V, Fernández-Urquiza M (2018) Explicit oral narrative intervention for students with Williams syndrome. Front Psychol. 8: 2337 http://doi.org/10.3389/fpsyg.2017.02337

Dunn L, Dunn LM, Arribas D (2006) Peabody, test de vocabulario en imágenes. Madrid: TEA Ediciones.

Endris V, Haussmann L, Buss E, Bacon C, Bartsch D, Rappold G. (2011) SrGAP3 interacts with lamellipodin at the cell membrane and regulates Rac-dependent cellular protrusions. J Cell Sci. 124(Pt 23):3941–55. doi: 10.1242/jcs.077081.

Endris V, Wogatzky B, Leimer U, Bartsch D, Zatyka M, Latif F, Maher ER, Tariverdian G, Kirsch S, Karch D, Rappold GA (2002) The novel Rho-GTPase activating gene MEGAP/ srGAP3 has a putative role in severe mental retardation. Proc Natl Acad Sci U S A. 99(18):11754–9. doi: 10.1073/pnas.162241099.

Engelhardt KR, Gertz ME, Keles S, Schäffer AA, Sigmund EC, Glocker C, Saghafi S, Pourpak Z, Ceja R, Sassi A, Graham LE, Massaad MJ, Mellouli F, Ben-Mustapha I, Khemiri M, Kilic SS, Etzioni A, Freeman AF, Thiel J, Schulze I, Al-Herz W, Metin A, Sanal Ö, Tezcan I, Yeganeh M, Niehues T, Dueckers G, Weinspach S, Patiroglu T, Unal E, Dasouki M, Yilmaz M, Genel F, Aytekin C, Kutukculer N, Somer A, Kilic M, Reisli I, Camcioglu Y, Gennery AR, Cant AJ, Jones A, Gaspar BH, Arkwright PD, Pietrogrande MC, Baz Z, Al-Tamemi S, Lougaris V, Lefranc G, Megarbane A, Boutros J, Galal N, Bejaoui M, Barbouche MR, Geha RS, Chatila TA, Grimbacher B (2015) The extended clinical phenotype of 64 patients with dedicator of cytokinesis 8 deficiency. J Allergy Clin Immunol 136: 402–12.

Farré A, Narbona J (2013). EDAH. Evaluación del Trastorno por Déficit de Atención con Hiperactividad. Madrid: TEA Ediciones, S. A.

Fernández-Urquiza M, Miranda M, Martínez V, Diez-Itza E (2016) Pragmática textual de las narraciones en el síndrome de Down: Perfiles de coherencia y cohesión. In E. Aguilar, D. Adrover, L. Bull, & R. López (Eds.), VIIIth International Conference of Language Acquisition (p. 30). Palma de Mallorca: UIB.

Fernández-Urquiza M, Viejo A, Cortiñas S, Huelmo J, Medina B, García I, Diez-Itza E (2015) Perfiles pragmáticos comparados de síndromes genéticos neuroevolutivos (S. Williams, S. Down y S. X Frágil). In F. Diéguez-Vide (Ed.), Temas de Lingüística Clínica (pp. 89–90). Barcelona: Horsori Editorial;

Fuchs S, Herzog D, Sumara G, Büchmann-Møller S, Civenni G, Wu X, Chrostek-Grashoff A, Suter U, Ricci R, Relvas JB, Brakebusch C, Sommer L (2009) Stage-specific control of neural crest stem cell proliferation by the small rho GTPases Cdc42 and Rac1. Cell Stem Cell. 2009 Mar 6;4(3):236–47. doi: 10.1016/j.stem.2009.01.017.

Garcia-Calero E, Botella-Lopez A, Bahamonde O, Perez-Balaguer A, Martínez S (2016) FoxP2 protein levels regulate cell morphology changes and migration patterns in the vertebrate developing telencephalon. Brain Struct. Funct. 221: 2905–2917.

Gilks WP, Hill M, Gill M, Donohoe G, Corvin AP, Morris DW (2012) Functional investigation of a schizophrenia GWAS signal at the CDC42 gene. World J. Biol. Psychiatry 13: 550–4.

Glessner JT, Li J, Wang D, March M, Lima L, Desai A, Hadley D, Kao C, Gur RE, Cohen N, Sleiman PMA, Li Q, Hakonarson H,; Janssen-CHOP Neuropsychiatric Genomics Working Group (2017) Copy number variation meta-analysis reveals a novel duplication at 9p24 associated with multiple neurodevelopmental disorders. Genome Med. 9: 106.

Gonçalves OF, Pinheiro AP, Sampaio A, Sousa N, Fernández M, Henriques M (2010) The narrative profile in Williams syndrome: There is more to storytelling than just telling a story. Br J Dev Disabil. 56(2): 89–109. doi:10.1179/096979510799102943

Gorelik A, Sapir T, Woodruff TM, Reiner O (2017) Serping1/C1 inhibitor affects cortical development in a cell autonomous and non-cell autonomous manner. Front Cell Neurosci. 11:169. doi: 10.3389/fncel.2017.00169.

Graham SA, Deriziotis P, Fisher SE (2015) Insights into the genetic foundations of human communication. Neuropsychol Rev. 25(1):3–26. doi: 10.1007/s11065-014-9277-2.

Griggs BL, Ladd S, Saul RA, DuPont BR, Srivastava AK (2008) Dedicator of cytokinesis 8 is disrupted in two patients with mental retardation and developmental disabilities. Genomics 91:195–202.

Hämmerle B, Carnicero A, Elizalde C, Ceron J, Martinez S, Tejedor FJ (2003) Expression patterns and subcellular localization of the Down syndrome candidate protein MNB/DYRK1A suggest a role in late neuronal differentiation. Eur J Neurosci. 17: 2277–86.

Hao HN, Zhao J, Lotoczky G, Grever WE, Lyman WD (2001) Induction of human beta-defensin-2 expression in human astrocytes by lipopolysaccharide and cytokines. J Neurochem. 77:1027–35.

Harada Y, Tanaka Y, Terasawa M, Pieczyk M, Habiro K, Katakai T, Hanawa-Suetsugu K, Kukimoto-Niino M, Nishizaki T, Shirouzu M, Duan X, Uruno T, Nishikimi A, Sanematsu F, Yokoyama S, Stein JV, Kinashi T, Fukui Y (2012) DOCK8 is a Cdc42 activator critical for interstitial dendritic cell migration during immune responses. Blood 119:4451–61.

Hernández A, Aguilar C, Paradell E, Valla F (2015) Revisión de la Adaptación Española de la Escala de Inteligencia de Wechsler para Niños - V. Madrid: Pearson Education

Hong SE, Shugart YY, Huang DT, Shahwan SA, Grant PE, Hourihane JO, Martin ND, Walsh CA (2000) Autosomal recessive lissencephaly with cerebellar hypoplasia is associated with human RELN mutations. Nat Genet. 26(1):93–6. doi: 10.1038/79246.

Hu SG, Zou M, Yao GX, Ma WB, Zhu QL, Li XQ, Chen ZJ, Sun Y (2014) Androgenic regulation of beta-defensins in the mouse epididymis. Reprod Biol Endocrinol. 12: 76.

Huang H, Tindall DJ (2007) Dynamic FoxO transcription factors. J Cell Sci. 120: 2479–87.

Infante J, Prieto C, Sierra M, Sánchez-Juan P, González-Aramburu I, Sánchez-Quintana C, Berciano J, Combarros O, Sainz J (2015) Identification of candidate genes for Parkinson’s disease through blood transcriptome analysis in LRRK2-G2019S carriers, idiopathic cases, and controls. Neurobiol Aging. 36(2):1105–9. doi: 10.1016/j.neurobiolaging.2014.10.039.

Irizarry RA, Hobbs B, Collin F, Beazer-Barclay YD, Antonellis KJ, Scherf U, Speed TP (2003) Exploration, normalization, and summaries of high density oligonucleotide array probe level data. Biostatistics 4: 249–264.

Ishiguro H, Hall FS, Horiuchi Y, Sakurai T, Hishimoto A, Grumet M, Uhl GR, Onaivi ES, Arinami T (2014) NrCAM-regulating neural systems and addiction-related behaviors. Addict Biol. 19(3):343–53. doi: 10.1111/j.1369-1600.2012.00469.x.

Ishiguro H, Miyake K, Tabata K, Mochizuki C, Sakurai T, Onaivi ES (2019) Neuronal cell adhesion molecule regulating neural systems underlying addiction. Neuropsychopharmacol Rep. 39(1):10–16. doi: 10.1002/npr2.12038.

Johnson-Glenberg MC (2008) Fragile X syndrome: Neural network models of sequencing and memory. Cogn Syst Res. 9:274–292. doi:10.1016/j.cogsys.2008.02.002

Juárez A, Monfort M (1998) Registro Fonológico Inducido. Madrid: CEPE.

Katayama K, Imai F, Campbell K, Lang RA, Zheng Y, Yoshida Y (2013) RhoA and Cdc42 are required in pre-migratory progenitors of the medial ganglionic eminence ventricular zone for proper cortical interneuron migration. Development. 140: 3139–45.

Kearney CJ, Randall KL, Oliaro J (2017) DOCK8 regulates signal transduction events to control immunity. Cell Mol Immunol 14: 406–411.

Krgovic D, Kokalj Vokac N, Zagorac A, Gregoric Kumperscak H (2018) Rare structural variants in the DOCK8 gene identified in a cohort of 439 patients with neurodevelopmental disorders. Sci Rep. 8: 9449.

Kurt S, Fisher SE, Ehret G (2012) Foxp2 mutations impair auditory-motor association learning. PLoS One 7: e33130.

Linzmeier RM, Ganz T (2005) Human defensin gene copy number polymorphisms: comprehensive analysis of independent variation in alpha- and beta-defensin regions at 8p22-p23. Genomics 86: 423–30.

Losh M, Bellugi U, Reilly J (2000) Narrative as a social engagement tool: The excessive use of evaluation in narratives from children with Williams syndrome. Narrat Inq. 10(2): 265–290. doi:10.1075/ni.10.2.01los

Ma Y, Mi Y-J, Dai Y-K, Fu H-L, Cui D-X, Jin WL (2013) The inverse F-BAR domain protein srGAP2 acts through srGAP3 to modulate neuronal differentiation and neurite outgrowth of mouse neuroblastoma cells. PLoS One 8, e57865.

MacWhinney B (2000) The CHILDES Project: Tools for Analyzing Talk. Volume I: Transcription format and programs. Mahwah: Lawrence Erlbaum.

Manga D, Ramos F (2006) Luria-inicial. Evaluación neuropsicológica en la edad preescolar. Madrid: TEA Ediciones.

Matzel LD, Babiarz J, Townsend DA, Grossman HC, Grumet M (2008) Neuronal cell adhesion molecule deletion induces a cognitive and behavioral phenotype reflective of impulsivity. Genes Brain Behav. 2008 Jun;7(4):470–80. doi: 10.1111/j.1601-183X.2007.00382.x.

Mendoza E, Carballo G, Muñoz J, Fresneda MD (2005) Test de Comprensión de Estructuras Gramaticales. Madrid: TEA.

Metsu S, Rooms L, Rainger J, Taylor MS, Bengani H, Wilson DI, Chilamakuri CS, Morrison H, Vandeweyer G, Reyniers E, Douglas E, Thompson G, Haan E, Gecz J, Fitzpatrick DR, Kooy RF (2014) FRA2A is a CGG repeat expansion associated with silencing of AFF3. PLoS Genet. 10(4):e1004242. doi: 10.1371/journal.pgen.1004242.

Miyamoto Y, Torii T, Kawahara K, Tanoue A, Yamauchi J (2016) Dock8 interacts with Nck1 in mediating Schwann cell precursor migration. Biochem Biophys Rep. 6:113–123.

Mohan V, Gomez JR, Maness PF (2019) Expression and function of neuron-glia-related cell adhesion molecule (NrCAM) in the amygdalar pathway. Front Cell Dev Biol. 7:9. doi: 10.3389/fcell.2019.00009.

Montero D (1996) Evaluación de la conducta adaptativa en personas con discapacidades. Adaptación y validación del ICAP. Bilbao: Mensajero.

Moore JM, Oliver PL, Finelli MJ, Lee S, Lickiss T, Molnár Z, Davies KE (2014) Laf4/Aff3, a gene involved in intellectual disability, is required for cellular migration in the mouse cerebral cortex. PLoS One 9(8):e105933. doi: 10.1371/journal.pone.0105933.

Moy SS, Nonneman RJ, Young NB, Demyanenko GP, Maness PF (2009) Impaired sociability and cognitive function in Nrcam-null mice. Behav Brain Res. 205(1):123–31. doi: 10.1016/j.bbr.2009.06.021.

Murphy E, Benítez-Burraco A (2016) Language deficits in schizophrenia and autism as related oscillatory connectomophathies: an evolutionary account. Neurosci. Biobehav Rev S0149-7634(16)30155-5.

Murphy E, Benítez-Burraco A (2017) Bridging the gap between genes and language deficits in schizophrenia: an oscillopathic approach. Front Hum Neurosci 10: 422.

Myers JP, Robles E, Ducharme-Smith A, Gómez TM (2012) Focal adhesion kinase modulates Cdc42 activity downstream of positive and negative axon guidance cues. J. Cell. Sci. 125, 2918–29.

Orlic-Milacic M, Kaufman L, Mikhailov A, Cheung AY, Mahmood H, Ellis J, Gianakopoulos PJ, Minassian BA, Vincent JB. (2014) Over-expression of either MECP2_e1 or MECP2_e2 in neuronally differentiated cells results in different patterns of gene expression. PLoS One. 9:e91742.

Otowa T, Yoshida E, Sugaya N, Yasuda S, Nishimura Y, Inoue K, Tochigi M, Umekage T, Miyagawa T, Nishida N, Tokunaga K, Tanii H, Sasaki T, Kaiya H, Okazaki Y (2009) Genome-wide association study of panic disorder in the Japanese population. J Hum Genet. 54(2):122–6. doi: 10.1038/jhg.2008.17.

Pagnamenta AT, Bacchelli E, de Jonge MV, Mirza G, Scerri TS, Minopoli F, Chiocchetti A, Ludwig KU, Hoffmann P, Paracchini S, Lowy E, Harold DH, Chapman JA, Klauck SM, Poustka F, Houben RH, Staal WG, Ophoff RA, O’Donovan MC, Williams J, Nöthen MM, Schulte-Körne G, Deloukas P, Ragoussis J, Bailey AJ, Maestrini E, Monaco AP; International Molecular Genetic Study Of Autism Consortium (2010) Characterization of a family with rare deletions in CNTNAP5 and DOCK4 suggests novel risk loci for autism and dyslexia. Biol Psychiatry. 68(4):320–8. doi: 10.1016/j.biopsych.2010.02.002.

Paracchini S, Diaz R, Stein J. (2016) Advances in dyslexia genetics—new insights into the role of brain asymmetries. In Friedmann, T., Dunlap, J. C., Goodwin, S. F. (Eds.), Advances in Genetics 96 (pp.53–97). London: Academic Press.

Pettigrew KA, Frinton E, Nudel R, Chan MTM, Thompson P, Hayiou-Thomas ME, Talcott JB, Stein J, Monaco AP, Hulme C, Snowling MJ, Newbury DF, Paracchini S. (2016) Further evidence for a parent-of-origin effect at the NOP9 locus on language-related phenotypes. J Neurodev Disord. 8, 24.

Pfenning AR, Hara E, Whitney O, Jarvis ED (2014) Convergent transcriptional specializations in the brains of humans and songlearning birds. Science 346(6215), 1256846. https://doi:10.1126/science.1256846.

Reilly J, Losh M, Bellugi U, Wulfeck B (2004) “Frog, where are you?” Narratives in children with specific language impairment, early focal brain injury, and Williams syndrome. Brain Lang 88(2): 229–247. doi:10.1016/S0093-934X(03)00101-9

Reinhard NR, Van Der Niet S, Chertkova A, Postma M, Hordijk PL, Gadella TWJ Jr, Goedhart J (2019) Identification of guanine nucleotide exchange factors that increase Cdc42 activity in primary human endothelial cells. Small GTPases. 30:1–15. doi: 10.1080/21541248.2019.1658509.

Ren M, Hu Z, Chen Q, Jaffe A, Li Y, Sadashivaiah V, Zhu S, Rajpurohit N, Heon Shin J, Xia W, Jia Y, Wu J, Lang Qin S, Li X, Zhu J, Tian Q, Paredes D, Zhang F, Wang KH, Mattay VS, Callicott JH, Berman KF, Weinberger DR, Yang F (2020) KCNH2-3.1 mediates aberrant complement activation and impaired hippocampal-medial prefrontal circuitry associated with working memory deficits. Mol Psychiatry. 25(1):206–229. doi: 10.1038/s41380-019-0530-1.

Reynolds CR, Kamphaus RW (2013) RIAS. Reynolds Intellectual Assessment Scales. Madrid: TEA Ediciones Ediciones, S. A.

Reynolds CR, Kamphaus RW (2004) Behavior Assessment System for Children-BASC-2. Madrid: TEA Ediciones

Rice ML, Smolik F, Perpich D, Thompson T, Rytting N, Blossom M (2010) Mean length of utterance levels in 6-month intervals for children 3 to 9 years with and without language impairments. J Speech Lang Hear Res. 53(2):333–49. doi: 10.1044/1092-4388(2009/08-0183).

Roid GH, Miller LJ, Pomplun M, Koch C (2013) LEITER – 3: Escala manipulativa international de Leiter (3^a^ edición). Madrid: Psymtec.

Sacks H, Schegloff E, Jefferson G (1974) A simplest systematics for the organization of turn-taking for conversation. Language 50(4): 696–735.

Sakurai T (2012) The role of NrCAM in neural development and disorders--beyond a simple glue in the brain. Mol Cell Neurosci. 49(3):351–63. doi: 10.1016/j.mcn.2011.12.002.

Sakurai T, Ramoz N, Reichert JG, Corwin TE, Kryzak L, Smith CJ, Silverman JM, Hollander E, Buxbaum JD (2006) Association analysis of the NrCAM gene in autism and in subsets of families with severe obsessive-compulsive or self-stimulatory behaviors. Psychiatr Genet. 16(6):251–7. doi: 10.1097/01.ypg.0000242196.81891.c9.

Schreiweis C, Bornschein U, Burguière E, Kerimoglu C, Schreiter S, Dannemann M, Goyal S, Rea E, French CA, Puliyadi R, Groszer M, Fisher SE, Mundry R, Winter C, Hevers W, Pääbo S, Enard W, Graybiel AM (2014) Humanized Foxp2 accelerates learning by enhancing transitions from declarative to procedural performance. Proc Natl Acad Sci U S A. 111(39):14253–8. doi: 10.1073/pnas.1414542111.

Semel E, Wiig EH, Secord WA (2006) Clinical Evaluation of Language Fundamentals. Spanish Edition (CELF-4). Barcelona: Pearson.

Shiro M, Diez-Itza E, Viejo A, Fernández-Urquiza M (2016) Pragmática evaluativa de las narraciones en el síndrome de Williams. In E. Aguilar, D. Adrover, L. Bull, & R. López (Eds.), Proceedings of the VIIIth International Conference of Language Acquisition (pp. 31–32). Palma de Mallorca: UIB.

Spiteri E, Konopka G, Coppola G, Bomar J, Oldham M, Ou J, Vernes SC, Fisher SE, Ren B, Geschwind DH (2007) Identification of the transcriptional targets of FOXP2, a gene linked to speech and language, in developing human brain. Am J Hum Genet. 81(6):1144–57. doi: 10.1086/522237.

Szklarczyk D, Franceschini A, Wyder S, Forslund K, Heller D, Huerta-Cepas J, Simonovic M, Roth A, Santos A, Tsafou KP, Kuhn M, Bork P, Jensen LJ, von Mering C. (2015) STRING v10: protein-protein interaction networks, integrated over the tree of life. Nucleic Acids Res. 43: D447–52.

Thurstone LL, Yela M (2012) CARAS-R. Test de Percepción de Diferencias-Revisado. Madrid: TEA Ediciones S.A.

Traylor M, Tozer DJ, Croall ID, Lisiecka-Ford DM, Olorunda AO, Boncoraglio G, Dichgans M, Lemmens R, Rosand J, Rost NS, Rothwell PM, Sudlow CLM, Thijs V, Rutten-Jacobs L, Markus HS; International Stroke Genetics Consortium (2019) Genetic variation in PLEKHG1 is associated with white matter hyperintensities (n = 11,226). Neurology 92(8):e749–e757. doi: 10.1212/WNL.0000000000006952.

Tsui D, Vessey JP, Tomita H, Kaplan DR, Miller FD (2013) Foxp2 regulates neurogenesis during embryonic cortical development. J Neurosci. 33: 244–258.

Turner SJ, Hildebrand MS, Block S, Damiano J, Fahey M, Reilly S, Bahlo M, Scheffer IE, Morgan AT (2013) Small intragenic deletion in FOXP2 associated with childhood apraxia of speech and dysarthria. Am J Med Genet A. 161A(9):2321–6. doi: 10.1002/ajmg.a.36055.

van Bon BW, Hoischen A, Hehir-Kwa J, de Brouwer AP, Ruivenkamp C, Gijsbers AC, Marcelis CL, de Leeuw N, Veltman JA, Brunner HG, de Vries BB (2011) Intragenic deletion in DYRK1A leads to mental retardation and primary microcephaly. Clin Genet. 79(3):296–9. doi: 10.1111/j.1399-0004.2010.01544.x. PMID: 21294719.

Vargha-Khadem F, Gadian DG, Copp A, Mishkin M (2005) FOXP2 and the neuroanatomy of speech and language. Nature Rev Neurosci. 6: 131–138.

Vernes SC, Oliver PL, Spiteri E, Lockstone HE, Puliyadi R, Taylor JM, Ho J, Mombereau C, Brewer A, Lowy E, Nicod J, Groszer M, Baban D, Sahgal N, Cazier JB, Ragoussis J, Davies KE, Geschwind DH, Fisher SE (2011) Foxp2 regulates gene networks implicated in neurite outgrowth in the developing brain. PLoS Genet. 7(7):e1002145. doi: 10.1371/journal.pgen.1002145.

Verschueren J (1999). Understanding pragmatics. London: Edward Arnold.

Waclaw RR, Ehrman LA, Merchan-Sala P, Kohli V, Nardini D, Campbell K (2017) Foxo1 is a downstream effector of Isl1 in direct pathway striatal projection neuron development within the embryonic mouse telencephalon. Mol Cell Neurosci. 80:44–51. doi: 10.1016/j.mcn.2017.02.003.

Waltereit R, Leimer U, von Bohlen Und Halbach O, Panke J, Hölter SM, Garrett L, Wittig K, Schneider M, Schmitt C, Calzada-Wack J, Neff F, Becker L, Prehn C, Kutscherjawy S, Endris V, Bacon C, Fuchs H, Gailus-Durner V, Berger S, Schönig K, Adamski J, Klopstock T, Esposito I, Wurst W, de Angelis MH, Rappold G, Wieland T, Bartsch D (2012) Srgap3^−^/^−^ mice present a neurodevelopmental disorder with schizophrenia-related intermediate phenotypes. FASEB J. 26(11):4418–28. doi: 10.1096/fj.11-202317.

Wang T, Guo H, Xiong B, Stessman HA, Wu H, Coe BP, Turner TN, Liu Y, Zhao W, Hoekzema K, Vives L, Xia L, Tang M, Ou J, Chen B, Shen Y, Xun G, Long M, Lin J, Kronenberg ZN, Peng Y, Bai T, Li H, Ke X, Hu Z, Zhao J, Zou X, Xia K, Eichler EE, (2016) De novo genic mutations among a Chinese autism spectrum disorder cohort. Nat Commun. 7:13316.

Wang Z, Hong Y, Zou L, Zhong R, Zhu B, Shen N, Chen W, Lou J, Ke J, Zhang T, Wang W, Miao X (2014) Reelin gene variants and risk of autism spectrum disorders: an integrated meta-analysis. Am J Med Genet B Neuropsychiatr Genet. 165B(2):192–200. doi: 10.1002/ajmg.b.32222.

Williams WM, Castellani RJ, Weinberg A, Perry G, Smith MA (2012) Do β-defensins and other antimicrobial peptides play a role in neuroimmune function and neurodegeneration? Sci. World J 2012: 905785.

Wilson NK, Lee Y, Long R, Hermetz K, Rudd MK, Miller R, Rapoport JL, Addington AM (2011) A novel microduplication in the neurodevelopmental gene SRGAP3 that segregates with psychotic illness in the family of a COS proband. Case Rep Genet. 2011:585893. doi: 10.1155/2011/585893.

Wong K, Ren XR, Huang YZ, Xie Y, Liu G, Saito H, Tang H, Wen L, Brady-Kalnay SM, Mei L, Wu JY, Xiong WC, Rao Y (2001) Signal transduction in neuronal migration: roles of GTPase activating proteins and the small GTPase Cdc42 in the Slit-Robo pathway. Cell. 107(2):209–21. doi: 10.1016/s0092-8674(01)00530-x. PMID: 11672528.

Xiang YY, Dong H, Wan Y, Li J, Yee A, Yang BB, Lu WY (2006) Versican G3 domain regulates neurite growth and synaptic transmission of hippocampal neurons by activation of epidermal growth factor receptor. J Biol Chem. 281(28):19358–68. doi: 10.1074/jbc.M512980200.

Yiin JJ, Hu B, Jarzynka MJ, Feng H, Liu KW, Wu JY, Ma HI, Cheng SY (2009) Slit2 inhibits glioma cell invasion in the brain by suppression of Cdc42 activity. Neuro Oncol. 11(6):779–89. doi: 10.1215/15228517-2008-017.

