## Supplemental file 2 for "Language impairment with a partial duplication of *DOCK8*"

@Begin

@Languages:	spa

@Participants:	CHI Informant, MOT Mother, SPL Speech Therapist, ADU Adult

@Options:	CA

@ID:	spa|CHROMOLANG|CHI|11;6.|male|||Informant|||

@ID:	spa|CHROMOLANG|MOT|||||Mother|||

@ID:	spa|CHROMOLANG|SPL|||||Therapist|||

@ID:	spa|CHROMOLANG|ADU|||||Adult|||

@Transcriber:	Maite Fernández Urquiza

@Time Duration:	00:15:15

@Date:	24-APR-2018

@Location:	Córdoba

@Situation:	Room at the Speech Language Therapy Clinic

@Activities:	Playing, talking about family and school

*MOT:	a ver@i (.) cuál <quieres [/-] qué color quieres>

[=! aspirated /s/, dialectal] ?

%sit:	CHI is searching for something into a box.

*CHI:	pero: falta el dojo [>][=! looking at SPL].

%err:	dojo=rojo $PHO

*SPL:	sí se: [<] [/-] está en casa (.) pero no pasa nada „ no?

*MOT:	bueno pues [=! aspirated /s/, dialectal] coge otro color (.) <qué te

gusta [=! aspirated /s/, dialectal] el rojo> [>]?

*SPL:	o qué [<] ?

*CHI:	no porque me gusta [=! aspirated /s/, dialectal] el dojo

[=! nods, smiles].

%err:	dojo=rojo $PHO

*CHI:	este [=! aspirated /s/, dialectal] me <xxx xxx>

[>][=! grabbing the blue elastic].

*SPL:	da igual [<].

*MOT:	el azul ?

*MOT:	vale@i yo cojo el verde [=! grabbing the green elastic].

%sit:	CHI puts the elastic around his head.

*MOT:	pero esto [=! aspirated /s/, dialectal] lo tienes

[=! aspirated /s/, dialectal] que <poner

[=! deleted /r/, dialectal] así> [>]

[=! tidying up the cards on the table].

*CHI:	sí sí sí [<] [=! helps with the tidying up].

*SPL:	explícaselo a mamá.

*CHI:	(0.5) e:h (0.4) tú: +/.

%xepr:	$et3:DRA:FIL

*SPL:	es al revés (.) pero bueno.

*ADU:	para arriba (.) que si no la tarjeta te va a tapar

[=! deleted /r/, dialectal] <los ojos> [=! deleted /s/, dialectal] .

*MOT:	tú la tienes [=! aspirated /s/, dialectal] bien

[=! looks at CHI and puts the elastic around his head upside down] ?

*ADU:	no (.) al revés [=! deleted /s/, dialectal] .

*CHI:	vale@i [=! tidying up the cards] yo primero.

*SPL:	pero no le has explicado a mamá cómo es el juego (.) ella no lo sabe

(.) explícaselo por favor.

*CHI:	que: [=! looking to MOT in the eyes] [/] <que tenemos> [/] que

tenemos [=! deleted /s/, dialectal] a [*] poner

[=! deleted /r/, dialectal] una tajeta [*] aquí

[=! holding one card over his forehead with the elastic] .

%err:	a=que $MOR $YN; tajeta=tarjeta $PHO;

%xepr:	$i5:MMN:REP; $i5:MMN:REP; $i5:MMN:ORD:SST; $i5:MQT:VUT

*MOT:	y ahí dónde es [=! deleted /s/, dialectal ] ?

*CHI:	aquí [=! holding the card over his forehead with the elastic again] .

%xepr:	$et4:RPR:OTH:PER

*CHI:	y: [/] <y yo:> [/] y yo digo: como: +"/.

%xepr:	$i5:MMN:REP; $i5:MMN:REP; $et3:DRA:FIL

*CHI:	+" es [=! aspirated /s/, dialectal] un animal ?

*CHI:	y: [/] y todos [=! deleted /s/, dialectal] [=! illustrator] dice:

[*] +"/.

%err:	dice=dicen $MOR $SYN

%xepr:	$i5:MMN:REP; $i5:MMN:ORD:SST

*CHI:	+" zí [*] (.) no .

%err:	zí=sí $PHO

*MOT:	vale@i [=! nods] .

*CHI:	y: así: <toro [*] el resto> [?] hum@i [=! nods].

%err:	toro=todo $PHO

%err:	$et3:DRA:FIL

*MOT:	entonces dividimos [=! deleted /s/, dialectal] [/-] adivinamos

[=! deleted /s/, dialectal] el tuyo primero

[=! pointing CHI with forefinger] ?

*CHI:	no [=! tidying up the cards].

*MOT:	yo hablo y tú me tienes [=! deleted /s/, dialectal] que: [/-] o cómo

lo hacemos [=! deleted /s/, dialectal] ?

*MOT:	me: [/] me pongo yo primero .

%sit:	CHI tries to hold a card on MOT's forehead with the elastic .

*CHI:	pimero [*] tú [=! smiles].

%err:	pimero=primero $PHO

*MOT:	vale@i yo empiezo.

*CHI:	espera [=! CHI is still trying to hold the card on MOT´s forehead].

*SPL:	no empieza tú primero porfa@d (.) que mamá te ponga uno que no hayas

visto va@i .

*CHI:	vale@i [=! looks embarrased at SPL].

*ADU:	sí porque ese l(o) has visto .

*SPL:	ese <l(o) ha visto> [>].

*MOT:	l(o) ha visto [<] ?

*MOT:	www.

*SPL:	www.

*MOT:	<espera que están> [=! aspirated /s/, dialectal] tod:as

[=! deleted /s/, dialectal] [=! searching a new card] +//.

*ADU:	tachá:n@i !

%sit:	MOT holds a new card on CHI's forehead.

*MOT:	bue:no@i !

*SPL:	andá@i [>] (.) qué chuli !

*ADU:	así es [<] emocionante „ eh@i ?

*CHI:	0 [=! laughs]. [+ trn]

*SPL:	<venga@i va@i> [!] <pregunta [/-] pregúntale a mamá> [>] .

*MOT:	hazme una pregunta a ver@i (.) que yo te

contesto [=! aspirated /s/, dialectal][<] .

*CHI:	e:s (.) [/] es u:n animal ?

%xepr:	$i5:MMN:REP

*MOT:	sí [=! nods].

*CHI:	o:h@i [=! smiles] e:s [/-] (.) hu:m@i tie:ne: [/] tiene cola?

%xepr:	$i5:MQT:RVP:INT; $i5:MMN:REF; $et3:DRA:FIL; $i5:MMN:REP

*MOT:	sí [=! nods] .

*CHI:	un león [=! looks MOT in the eyes smiling].

%xepr:	$i5:MQL:QST:WOR:IMM

*MOT:	un qué ?

*CHI:	un león [=! shouting] !

%xepr:	$et4:RPR:OTH

*MOT:	no [=! shakes head no] .

*CHI:	hi:@i [=! shakes head ].

%xepr:	$i5:MQT:RVP:INT

*MOT:	ahora tú a mí .

*CHI:	e:h +/.

%xepr:	$et3:DRA:FIL

*MOT:	no no espérate [=! aspirated /s/, dialectal] yo a ti !

*ADU:	pregunta .

*MOT:	e:s [=! deleted /s/, dialectal] un animal ?

*CHI:	no [=! shakes head no] .

%xepr:	$et3:GSA

*MOT:	es [=! deleted /s/, dialectal] una cosa ?

*CHI:	zí [*][=! nods] .

%err:	zí=sí $PHO

%xepr:	$et3:GSA

*MOT:	<e:s [/] es> [=! deleted /s/, dialectal] u:n mueble ?

*CHI:	no [=! smiles].

*MOT:	e:s [=! deleted /s/, dialectal] u:n aparato?

*CHI:	no [=! shakes head no] .

*SPL:	bueno@i ahora él venga@i [>].

*MOT:	es un juego [<] ?

*CHI:	0 [=! laughs] . [+ trn]

*MOT:	0 [=! laughs] . [+ trn]

*SPL:	ahora él .

*ADU:	venga@i te toca preguntar a ti [=! to CHI].

*SPL:	venga@i.

*CHI:	tengo: [/] tengo: [/-] qué: [*] color zoy [*] ?

%err:	qué=de qué $MOR $SYN zoy=soy $PHO

%xepr:	$i5:MMN:REP; $i5:MMN:REF; $i5:MMN:ORD:OMI

*MOT:	de qué color eres [=! deleted /s/, dialectal] ?

*MOT:	amarillo y con <pintitas marrones> [=! deleted /s/, dialectal] .

*CHI:	e:h buf@i [=! looks away while thinking] (.) un perro

[=! looks at MOT smiling].

%xepr:	$et3:DRA:FIL; $i5:MQT:RVP:INT; $i5:PIM:PMQT:ADQ; $i5:MQL:QST:WOR:IMM

*MOT:	un perro no (.) e:h sirve para botar [?] es

[=! deleted /s/, dialectal] un objeto [/-] un juguete ?

*CHI:	no [=! shakes head].

%xepr:	$et3:GSA

*MOT:	xxx xxx xxx se me cae [=! laughs] .

%sit:	MOT is fixing the elastic around her head.

*CHI:	0 [=! laughs] .

*MOT:	e:h sirve para guardar

[=! deleted /r/ in final position of the word, dialectal] cosas

[=! deleted /s/ in final position of the word, dialectal] ?

*CHI:	no [=! shakes head no] .

%xepr:	$et3:GSA

*MOT:	hu:m@i sirve para: (.) para qué sirve [=! laughs] ?

%sit:	everybody laughs

*ADU:	a ver@i dale tú una pista [=! aspirated /s/ dialectal].

*MOT:	para qué sirve dímelo que no lo sé [=! laughing] .

*SPL:	dale una pista venga@i.

*CHI:	e:s [=! deleted /s/, dialectal] una fruta .

*MOT:	es [=! deleted /s/, dialectal] una [/-] ah@i es

[=! deleted /s/, dialectal] una fruta !

*CHI:	0 [=! nods, laughs]. [+ trn]

%xepr:	$et3:GSA

*MOT:	<está [=! aspirated /s/, dialectal] dulce> [/-] <es dul> [/-] es

dulce [=! aspirated /s/, dialectal] ?

*CHI:	e:h zí [*] [=! nods] .

%xepr:	$et3:DRA:FIL; $et3:GSA

%err:	zí=sí $PHO

*MOT:	u:na manzana .

*CHI:	no .

*MOT:	vaya por dios@i [=! deleted /s/, dialectal] (.) a ver@i

[=! deleted /r/, dialectal] tú a mí .

*CHI:	(0.4) tengo: [/-] soy un conejo ?

%xepr:	$i5:MMN:REF; $i5:MQL:QST:WOR

*MOT:	hum@i pero pregunta cosas [=! deleted /s/, dialectal] para:

adivinarlo .

*CHI:	no lo sé: [=! hand over his mouth] .

%xepr:	$et3:DRA:ERQ

*MOT:	pregunta más cosas si tiene: [/-] <cuántas patas y to(d)as@d esas

cosas> [=! deleted /s/ in final position of the word, dialectal] [>]

.

*CHI:	ah@i <cuántas [<] patas> [=! deleted /s/, dialectal] tiene

[=! touches MOT's arm to get her attention]?

%xepr:	$i5:MQT:RVP:INT; $i5:MQT:RUT:ECO

*MOT:	cuatro [=! laughs] .

*CHI:	(0.4) no sé [=! smiles] a:a [*] tú

[=! touches MOT's arm to give her the floor].

%err:	a:a=ahora $PHO

%xepr:	$et3:DRA:ERQ

*MOT:	e:h es [=! deleted /s/, dialectal] de

color [=! deleted /r/, dialectal] amarillo ?

*CHI:	no [=! shakes head no] .

%xepr:	$et3:GSA; $i5:MQL:ASW:IMM

*MOT:	de color rojo?

*CHI:	no [=! shakes head no] .

%xepr:	$et3:GSA

*MOT:	de colo:r [=! deleted /r/, dialectal] +/.

*SPL:	de color amarillo no:?

*CHI:	ah@i <zí zí zí> [>] .

%xepr:	$et4:RPR:OTH; $i5:MQT:RVP:INT

*MOT:	de color amarillo un plátano [<] ?

*SPL:	algo de amarillo tiene „ no ?

*MOT:	un plátano ?

*CHI:	no .

*MOT:	vaya por dios@i [=! deleted /s/, dialectal].

*SPL:	tanto amarillo no .

*MOT:	tanto amarillo no?

*CHI:	no .

*MOT:	tiene rabillo así [=! illustrator with hand] ?

*CHI:	no [=! coming closer to MOT and looking her in the eyes, smiling].

*MOT:	<xxx fruta xxx xxx> [>] .

*SPL:	tú <has comido alguna vez> [/] has comido alguna vez esa fruta ?

*CHI:	zí [*] [=! nods, smiles].

%err:	zí=sí $PHO

%xepr:	$et3:GSA

*SPL:	y te gusta ?

*CHI:	zí [*] [=! nods looking at SPL].

%err:	zí=sí $PHO

%xepr:	$et3:GSA

*MOT:	yo llevo mucho a casa esa fruta ?

*CHI:	zí [*] [=! laughs looking at MOT].

%err:	zí=sí $PHO

*MOT:	y es [=! deleted /s/, dialectal] amarilla ?

*CHI:	zí [*] .

%err:	zí=sí $PHO

*MOT:	(0.5) una ciruela [=! illustrator with hand, laughs].

*CHI:	no [=! laughs] .

*ADU:	xxx xxx xxx tiene otro color.

*MOT:	ah@i que tiene otro color !

*MOT:	que no es [=! deleted /s/, dialectal] ama [/-] ah@i pero cuando

madura tiene otro color ?

*CHI:	zí [*] .

%err:	zí=sí $PHO

%xepr:	$i5:MQL:ASW:WOR

*MOT:	amarilla es [=! deleted /s/, dialectal] <antes

[=! deleted /s/, dialectal] de madurar

[=! deleted /r/ in final position of the word, dialectal] o cómo es

[=! deleted /s/, dialectal] eso> [>]?

*ADU:	a ver@i díselo tú xxx xxx xxx xxx [<] .

*CHI:	e:h te doy una pizta [*] o no [=! staring at MOT] ?

%err:	pizta=pista $PHO

%xepr:	$et3:DRA:FIL

*MOT:	vale@i dime [/-] dámela dámela .

*CHI:	za [/-] e:s [=! leans back, laughs, hand over mouth] e:s de: hum@i

+/.

%xepr:	$i5:MMN:REF; $et3:DRA:FIL

*ADU:	dale la pista de <los colores> [=! deleted /s/, dialectal] que tiene

.

*MOT:	a ver@i dime <los colores> [=! deleted /s/, dialectal] (.) que yo me

haga una idea.

*CHI:	tie:ne +/.

*MOT:	a ver@i dime la forma que tiene .

*CHI:	tie:ne +/.

*MOT:	qué forma tiene .

*CHI:	como: hum@i [=! shape illustrator with hand] así .

%xepr:	$et3:DRA:FIL; $et3:DRA:FIL; $et3:GSA: $et3:MQT:VUT

*MOT:	ese [/-] esa qué forma [/-] cuál es ?

*CHI:	triángula [*].

%err:	triángula=triangular $MOR

%xepr:	$i5:MMN:ORD:SST

*MOT:	triangular tiene forma triangular ah@i .

*MOT:	y tiene pepitas [=! deleted /s/, dialectal] ?

*CHI:	zí [*] .

%err:	zí=sí $PHO

%xepr:	$i5:MQL:ASW:WOR

*MOT:	0 [=! thinking].

*CHI:	zí [*] [=! looking at MOT in the eyes].

%err:	zí=sí $PHO

*MOT:	es un melón ?

*CHI:	hum@i casi [=! nods] !

%xepr:	$et3:DRA:FIL; $et3:GSA; $i5:MQL:ASW:WOR

*MOT:	casi uy@i [=! laughs, illustrator] m(e) ha faltao@d poco.

*CHI:	casi casi [=! nods].

%xepr:	$et3:GSA

*MOT:	te gusta [=! aspirated /s/, dialectal] mucho ?

*MOT:	en verano se come esa fruta ?

*CHI:	sí oy@i [=! leans back] .

%sit:	there is a fly in the room and CHI tries to get away, everybody

laughs.

*MOT:	bueno@i dime otra pista [=! aspirated /s/, dialectal] y ahora

seguimos [=! deleted /s/, dialectal] contigo

[=! taps CHI on the back].

*CHI:	e:s +/.

*MOT:	es [=! deleted /s/, dialectal] dulce o amarga ?

*CHI:	dulce [=! laughs] .

%sit:	the fly is annoying everybody in the room, CHI starts laughing,

his card falls down and he manages to see it's picture when picking

it up.

*MOT:	www.

*ADU:	www.

*CHI:	<ya zé> [*] [/] ya zé qué es [=! deleted /s/, dialectal]

%err:	zé=sé $PHO

%xepr:	$i5:MMN:REP

*MOT:	qué es [=! deleted /s/, dialectal] ?

*CHI:	u:n [/-] una jirafa [=! smiling] .

%xepr:	$i5:MMN:REF

*MOT:	porque l(a) <has visto> [=! aspirated /s/, dialectal] [=! lauhgs].

*ADU:	ay@i l(a) <has visto> [=! aspirated /s/, dialectal] !

%sit:	everybody laughs and the game comes to the end.

*MOT:	www.

*CHI:	www.

*ADU:	www.

*SPL:	www.

*MOT:	bueno@i <vamos a contarles [=! assimilation /r/>/l/, dialectal] a

ellas> [=! deleted /s/, dialectal] qué ha pasado hoy con tu tío (.)

qué le ha pasa(d)o@d al tío ?

*SPL:	andá@i !

*CHI:	que: hoy [/] <hoy mi tío> [/] hoy mi tío Enrique ze [*] opera .

%err:	ze=se $PHO

%xepr:	$i5:MMN:REP; $i5:MMN:REP

*MOT:	<y de qué se opera „ Joaquín> [>] ?

*SPL:	andá@i [<] !

*ADU:	xxx xxx [<] .

*CHI:	de: [/] de: <creo: de:> [/-] no sé [=! looks at MOT] .

%xepr:	$i5:MMN:REP; $i5:MMN:REF; $et3:DRA:ERQ

*MOT:	una hernia: +//.

*CHI:	de una henia [*] .

%err:	henia=hernia $PHO

*MOT:	y dónde está la hernia (.) en qué parte del cuerpo?

*CHI:	0 [=! touching his low thorax] . [+ trn]

%xepr:	$et3:GSA

*MOT:	un poquillo más [=! deleted /s/, dialectal] abajo [=! laughs] .

*SPL:	en la barriga [>] ?

*ADU:	vaya@i [<].

*MOT:	y de: [<] [/-] y tu tío cómo se llama ?

*CHI:	Anrique [*] .

%err:	Anrique=Enrique $PHO

*MOT:	Anrique ?

*CHI:	Enrique .

%xepr:	$et4:RPR:OTH

*MOT:	eso sí [=! nods] y de quién es [=! deleted /s/, dialectal] hermano ?

*CHI:	de mi padre .

*MOT:	y <cuántos hermanos> [=! deleted /s/, dialectal] tiene papá ?

*CHI:	prff@i [=! looks away] .

%xepr:	$i5:MQT:RVP:INT; $et3:GSA; $i5:PIM:PMQT:ADQ; $i5:MQL:ASW:WOR

*ADU:	sí tantos [=! aspirated /s/, dialectal] tiene ?

*MOT:	0 [=! looks to ADU indicating that the answer is not right].

*SPL:	qué suerte !

*MOT:	<cuántos hermanos> [=! deleted /s/, dialectal] tiene papá

[=! showing EBM] ?

*CHI:	muchos hum@i [=! looking MOT in the eyes] ?

%xepr:	$et3:DRA:FIL; $i5:MQL:QST:WOR; $et4:RPR:OTH:PER

*MOT:	solo tiene él [=! shakes head no, one EBM, laughs] <xxx xxx> [>].

*SPL:	andá@i [<] !

*CHI:	<ah@i va> [>] +/.

%xepr:	$i5:MQT:RVP:INT

*MOT:	<los demás> [=! deleted /s/, dialectal] son [/-] <los demás>

[=! deleted /s/, dialectal] qué son ?

*CHI:	ah@i vale@i primos [=! deleted /s/, dialectal] !

%xepr:	$i5:MQT:RVP:INT; $et4:RPR:OTH

*MOT:	claro@i <los demás son primos>

[=! deleted /s/ in final position of the word, dialectal] !

*CHI:	ah@i ez [*] <que creía que son [*] hermanos> [>].

%err:	ez=es $PHO son=eran $MOR $SYN

*MOT:	xxx xxx se llevan mu@d bien xxx xxx [<].

*ADU:	no es lo mismo primo (.) que hermano .

*SPL:	y la mamá cuántos hermanos tiene (.) la mamá ?

*CHI:	(0.4) muchos [=! smiles] .

*MOT:	eso sí [=! nods] .

*SPL:	ves ?

*MOT:	yo sí tengo muchos [=! deleted /s/, dialectal] .

*SPL:	muy bien (.) <cuántos tiene „ a ver@i> [>] ?

*MOT:	cuántos tengo [<] ?

*CHI:	(0.4) no: sé: [=! shakes head no] e:h una dos tres [=! whispered]

cuatro creo [=! frowns looking at MOT] .

%xepr:	$et3:DRA:ERQ; $et3:GSA; $et3:DRA:FIL;

*MOT:	cinco niñas [=! deleted /s/, dialectal] <y yo> [/-] y: bueno y

un niño.

*CHI:	el tío [=! fricative /t/] Pepe .

*MOT:	claro@i el único macho .

*CHI:	y: buf@i +/.

%xepr:	$i5:MQT:RVP:INT

*SPL:	y tú qué ?

*SPL:	tú tienes hermanos ?

*CHI:	zí [*] dos .

%err:	zí=sí $PHO

*SPL:	pero hermanos o hermanas ?

*CHI:	hermanas [=! deleted /s/, dialectal] .

*MOT:	dile dónde está [=! aspirated /s/, dialectal] Laura ahora .

*CHI:	en Alemania [=! troubled articulation].

*MOT:	pero <ahora dónde se ha ido> [/-] ahora dónde se ha ido Laura de

viaje ?

*CHI:	a: no sé .

%xepr:	$et3:DRA:ERQ

*MOT:	que ha ido a [/-] una semana de vacaciones a lo:s +//.

*CHI:	++ a los [=! deleted /s/, dialectal] An +/.

*MOT:	A:l +//.

*CHI:	++ A:lpes .

%xepr:	$et4:RPR:OTH

*MOT:	a <los Alpes> [=! deleted /s/, dialectal] [=! nods] .

*ADU:	pos@d qué suerte tiene tu hermana !

*SPL:	uhum@i en los Alpes qué hace frío o calor ?

*CHI:	frío [=! stands up and sits down again].

*MOT:	y qué s(e)@d ha lleva(d)o@d ?

*CHI:	a Toni [=! shrugs, laughs] .

%sit:	everybody laughs

%xepr:	$i5:MRL:ASW:TNG:ICOM

*MOT:	hombre@i ya@i [=! showing EBM] al novio pero aparte <de Tony qué

tiene> [>] [/] qué tiene más .

*ADU:	lo fundamental [<] .

*CHI:	el [/-] un [/-] dos perras [=! licking illustrator].

%xepr:	$i5:MMN:REF; $i5:MMN:REF

*ADU:	y qué hace tu hermana en Alemania ?

*CHI:	tabajar [*] [=! deleted /r/ with open /a/, dialectal] .

%err:	tabajá=trabajar $PHO

*ADU:	trabaja en Alemania?

*CHI:	0 [=! nods] . [+ trn]

%xepr:	$et3:GSA

*ADU:	y por qué se ha ido tan lejos [=! deleted /s/, dialectal] ?

*CHI:	po:s@d [=! deleted /s/, dialectal] para: [/-] (.) <para que:> [/]

pala [*] que: <pued(a) i(r) a> [?] tabajá [*]

[=! deleted /r/ with open /a/, dialectal] xxx .

%err:	pala=para $PHO; tabajá=trabajar $PHO

%xepr:	$i5:MMN:REF; $i5:MMN:REP; $i5:MQT:RUT; $i5:MQT:EVP

*MOT:	quería trabajar [=! deleted /r/, dialectal] „ no ?

*CHI:	quería trabajar [=! deleted /r/, dialectal] fuera .

%xepr:	$et4:RPR:OTH

*ADU:	y tu hermana en qué trabaja ?

*CHI:	o:h no sé [=! looks to MOT searching for help] .

%xepr:	$et3:DRA:FIL; $et3:DRA:ERQ; $et3:GSA

*MOT:	inter [/-] es [=! deleted /s/, dialectal] intérprete de: [/-] en

una: academia (.) está [=! aspirated /s/, dialectal] haciendo: [/-]

traduciendo en españo:l [=! aspirated /s/, dialectal] y en alemán .

*CHI:	vaya@i [=! nods] .

%xepr:	$et3:GSA; $i5:MQT:RVP:INT

*MOT:	y en <inglés tiene los tre:s> [=! deleted /s/, dialectal] .

*SPL:	muy bien (.) y a ti qué es lo que más te gusta hacer ?

*SPL:	a ver@i diles a ellas que no lo saben <yo sí lo sé> [>].

*ADU:	eso cuéntanos [=! deleted /s/, dialectal] lo que te gusta

[=! aspirated /s/, dialectal] [<] .

*CHI:	yo: [*] ziempre [*] me guta [*] jugar [=! deleted /r/, dialectal]

to:d [/-] cazi [*] to(dos)@d <los días> [=! deleted /s/, dialectal]

a la Play .

%err:	yo=a mí $MOR $SYN; ziempre=siempre $PHO; guta=gusta $PHO; cazi=casi

$PHO

%xepr:	$i5:MMN:ORD:OMI; $i5:MMN:ORD:SST; $i5:MMN:REF

*ADU:	anda@i <qué xxx> [>].

*MOT:	pero qué pasa qué te: [<] [/-] mamá qué te dice ?

*CHI:	que te [/-] me ve y me [?] dice +"/.

%xepr:	$i5:MMN:REF;

*CHI:	+" que no: (.) que no: !

*MOT:	que hay que hacer [=! deleted /r/, dialectal] qué ?

*CHI:	<las taleas> [=! deleted /s/, dialectal] [*] [=! laughs].

%err:	taleas=tareas $PHO

%sit:	everybody laughs.

*ADU:	te mandan <muchas tareas> [=! deleted /s/, dialectal] ?

*SPL:	pero aparte de la Play +//.

*MOT:	<martirio chino> [>] [=! illustrator with hand].

*SPL:	ah@i bueno@i perdona [<] .

*SPL:	te mandan muchas tareas dice Encarna?

*CHI:	zí [=! looks to SPL, then to ADU, nods] <unas pocas [?]>

[=! dialectal aspirated /s/, whispered, nods] .

%err:	zí=sí $PHO

%xepr:	$et3:GSA

*MOT:	sí pero e:h <qué te> [/-] qué m(e) has [=! dialectal aspirated /s/]

dicho hoy que te mandan <tareas de algo que no has>

[=! deleted /s/, dialectal] visto [=! aspirated /s/, dialectal] „ no

?

*CHI:	sí [=! nods] .

%xepr:	$et3:GSA

*SPL:	que es qué ?

*MOT:	que te tengo que explicar [=! deleted /r/, dialectal] yo y me

enfado [>] digo +"/.

*MOT:	+" <esto lo has> [=! aspirated /s/] tenido que ver

[=! deleted /r/, dialectal].

*ADU:	xxx xxx xxx xxx [<].

*SPL:	cómo que no has visto [<] ?

*MOT:	y tú qué me dices [=! deleted /s/, dialectal] ?

*CHI:	(0.4) hum [=! looks to MOT, then to ADU searching for help, smiles] .

%xepr:	$et3:GSA; $et3:DRA:FIL

*ADU:	que no lo <has visto> [=! dialectal aspirated /s/] .

*CHI:	no [=! looks to MOT, smiles].

*ADU:	a ver@i cuál era la tarea cuéntanos [=! dialectal aspirated /s/] .

*CHI:	e:la [*] [/] ela [*] poner [=! deleted /r/, dialectal] xxx xxx

poniendo uno:s [/] <unos eulos [*]> [=! deleted /s/, dialectal] y

otro [*] eulos [*] .

%err:	ela=era $PHO ela=era $PHO eulos=euros $PHO otro=otros $MOR

%xepr:	$i5:MMN:REP; $i5:MMN:REP; $i5:MMN:ORD:OMI; $i5:MQT:VUT

*CHI:	y: [/] <y: pa:(ra)@d> [/] y pa:@d hacer xxx xxx <y había> [/] y

<había como:> [/] había co:mo: [/] como <billetes y eulos [*]>

[=! deleted /s/, dialectal] y ya a:h debajo

[=! deictic illustrator] billetes [=! deleted /s/, dialectal] y

euros .

%err:	eulos=euros $PHO

%xepr:	$i5:MMN:REP; $i5:MMN:REP; $i5:MMN:REP; $i5:MMN:REP;

$et3:DRA:FIL; $i5:MMN:REP; $et3:DRA:FIL; $i5:MQT:RUT; $i5:MQT:EVP

*MOT:	pero qué tenías [=! deleted /s/, dialectal] que hacer con el

ejercicio ?

*CHI:	una suma creo .

%xepr:	$et4:RPR:OTH

*MOT:	contarlo [=! assimilation /r/ > /l/, dialectal] „ no

[=! showing EBM] ?

*CHI:	contarlo [=! assimilation /r/ > /l/, dialectal, nods] .

%xepr:	$i5:MQT:RUT:ECO

*MOT:	contar el dinero.

*SPL:	muy bien [>].

*ADU:	ah@i [<].

*MOT:	y luego de lengua qué te ha mandado ?

*CHI:	ajedtivos [*] .

%err:	ajedtivos=adjetivos $PHO

*SPL:	uhum@i.

*MOT:	y qué más ?

*CHI:	y: +/.

*MOT:	señalar [=! deleted /r/, dialectal] [<] qué ?

*CHI:	+, y señalar [=! deleted /r/, dialectal] <los velbos[*]>

[=! deleted /s/, dialectal] .

%err:	velbos=verbos $PHO

*MOT:	no (.) verbos [=! deleted /s/, dialectal] no [=! shakes head no].

*MOT:	la síla:(ba) +//.

*CHI:	++ la zila .

%err:	zila=sílaba $PHO

%xepr:	$i5:MQT:RUT:ECO:ICOM

*MOT:	la sílaba qué ?

*CHI:	++ tónica .

*MOT:	y: (.) luego la:s +//.

*CHI:	y: a ver@i <que no:> [/-] [=! whispers, effortful facial expression]

que: [/] que no me acueldo [*] [=! smiles, looks to MOT in the eyes].

%err:	acueldo=acuerdo $PHO

%xepr:	$et3:DRA:FIL; $i5:MMN:REF; $i5:MMN:REP; $et3:DRA:ERQ; $et3:GSA

*MOT:	<las agudas y las> [=! deleted /s/, dialectal] +//.

*CHI:	++ <las dicimales [*]> [=! deleted /s/, dialectal] .

%err:	dicimales=decimales $PHO

%xepr:	$i5:MRL:ASW:NRL

*MOT:	<las agudas y las> [=! deleted /s/, dialectal] +//?

*CHI:	++ y las [=! looking at MOT] ?

%xepr:	$i5:MQT:RVP:ECO; $et3:GSA

*MOT:	lla: +//.

*CHI:	++ llanas [=! deleted /s/, dialectal] .

*MOT:	<las agudas y las llanas> [=! deleted /s/, dialectal] .

*MOT:	y <les explicas> [=! deleted /s/, dialectal] [=! showing EBM] qué

<estás estudiando en naturales>

[=! aspirated /s/ in postnuclear position; deleted /s/ in final position

of the word] ?

*MOT:	<es que ellas [=! pointing to ADU and SPL] son profesoras>

[=! deleted /s/in final position of the word, dialectal] .

*ADU:	(cl)aro@i !

*CHI:	<pes [*] yo:> [/-] yo pes [*] yo <ahora <estoy estudiando>> [=!

aspirated /s/, dialectal] [/] ahora <estoy estudiando>

[=! aspirated /s/, dialectal] [=! looks to MOT] hum@i +/.

%err:	pes=pues $PHO

%xepr:	$i5:MMN:REF; $i5:MMN:REP; $et3:DRA:FIL

*ADU:	a ver .

*CHI:	0 [=! regulator: forefinger over his mouth to indicate ADU to remain silent].[+ trn]

%xepr:	$et3:GSA

*CHI:	+, <toy tudiando> [*] el [/] el cuelpo [*] humano .

%err:	toy tudiando=estoy estudiando $PHO cuelpo=cuerpo $PHO

%xepr:	$i5:MMN:REP

*SPL:	ah@i <qué bien> [>].

*ADU:	qué interesante [<] !

*MOT:	y qué funciones [=! deleted /s/, dialectal] tiene el cuerpo humano ?

*MOT:	<funciones vitales (.) cuáles> [=! deleted /s/, dialectal] son „

Joaquín ?

*CHI:	e:h tie: [/-] zo:n [*] [/-] ay@i que no me acuerdo .

%err:	zon=son $PHO

%xepr:	$et3:DRA:FIL; $i5:MMN:REF; $i5:MMN:REF; $et3:DRA:ERQ

*MOT:	son tres [=! deleted /s/, dialectal] [=! three EBM] xxx xxx xxx xxx

xxx [=! laughs] .

*MOT:	son <tres cuáles> [=! deleted /s/, dialectal] son <las funciones

vitales> [=! deleted /s/, dialectal] [=! laughs, three EBM] ?

*CHI:	0 [=! looks to MOT and laughs] . [ + trn ]

*ADU:	a ver@i <cuéntanos lo que tú te acuerdes>

[=! deleted /s/, dialectal] .

*CHI:	ps@i nada [=! shrugs] .

%xepr:	$i5:MQT:RVP:INT; $et3:GSA

*ADU:	es [=! deleted /s/, dialectal] que a mí se me han olvida(d)o@d <las

funciones vitales> [=! deleted /s/, dialectal] .

*SPL:	y te acuerdas qué has hecho hoy en el cole(gio)?

*CHI:	hoy sí .

*SPL:	a ver cuéntanos cuando llegaste esta mañana qué te pusieron ?

*CHI:	hoy: me: [/-] hoy he reza(d)o@d (.) después

[=! dialectal aspirated /s/] me he ido a:l [/-] &ao a otra claze [*]

.

%err:	claze=clase $PHO

%xepr:	$i5:MMN:REF; $i5:MMN:REF; $et3:DRA:DIS

*CHI:	después [=! aspirated /s/ in postnuclear position, deleted /s/ in final

position of the word, dialectal] venía [*] a mi claze [*] +/.

%err:	venía=he ido $MOR claze=clase $PHO

%xepr:	$i5:MMN:ORD:SST;

*ADU:	pero a qué te has [=! deleted /s/, dialectal] ido a otra clase ?

*ADU:	allí qué has [=! deleted /s/, dialectal] dado ?

*CHI:	apoyo .

*ADU:	sí pero qué (.) qué has [=! deleted /s/, dialectal] dado ?

*MOT:	qué has [=! deleted /s/, dialectal] dado de materia ?

*CHI:	lengua y mates siempre .

*ADU:	eso [?] lengua y mates [=! whispered].

*CHI:	y después [=! aspirated /s/ in postnuclear position, deleted /s/ in final

position of the word, dialectal] in +/.

*MOT:	y qué te dio ayer [=! deleted /r/, dialectal] la seño(rita) porque te

habías [=! deleted /s/, dialectal] portado bien ?

*CHI:	u:n [/-] dos [=! two EBM] potte [*][=! deleted /r/, dialectal] del

Real+Madrí@d y: una libreta.

%err:	potte=póster $PHO

%xepr:	$i5:MMN:REF; $et3:GSA;

*ADU:	bueno@i qué suerte [!] tú eres [=! deleted /s/, dialectal] del

Madrid „ Joaquín ?

*CHI:	claro@i [=! shrugs, smiles] !

*ADU:	igual que yo (.) de <los buenos> [=! deleted /s/, dialectal] „

verdad ?

*CHI:	claro@i [=! nods] !

%xepr:	$et3:GSA

*ADU:	verás [=! deleted /s/, dialectal] mañana (.) mañana vamos

[=! deleted /s/, dialectal] a ganá [=! deleted /r/, dialectal] al

xxx .

*MOT:	y luego que has [=! deleted /s/, dialectal] hecho ?

*CHI:	después [=! aspirated /s/ in postnuclear position, deleted /s/ in final

position of the word, dialectal] <me he ido> [=! syllabified] a:

[/] a ve:r religión (.) después +/.

%xepr:	$i5:MMN:REP

*MOT:	cómo se llama el profesor ?

*CHI:	Don+Juan .

*MOT:	y qué le pasa a ese profesor [=! looks to CHI and laughs] ?

*CHI:	que: [/] [=! looks to MOT and laughs] que rita [*] mucho [=! nods].

%err:	rita=grita $PHO

%sit:	everybody laughs

%xepr:	$i5:MMN:REP; $et3:GSA

*SPL:	y en religión qué os explican ?

*SPL:	de qué os hablan en religión?

*CHI:	de na@d es [=! deleted /s/, dialectal] que mientras

[=! deleted /s/, dialectal] él habla to:(dos)@d (es)tamos [*]

[=! deleted /s/, dialectal] charlando

[=! assimilation /r/ > /l/, dialectal] [=! shrugs].

%err:	tamos=estamos $PHO

%xepr:	$et3:DRA:FIL

*SPL:	ah@i él habla y vosotros +/.

*MOT:	pues por eso grita porque <no le hacéis [=! deleted /s/, dialectal]

caso> [>].

*SPL:	ah@i claro@i hombre:@i [<] .

*CHI:	hacía asto [*] +"/.

%err:	asto=esto $PHO

*CHI:	+" bla bla bla bla bla bla bla bla.

%xepr:	$i5:MQT:VUT

*CHI:	y Don+Juan como: +"/.

*CHI:	+" palalos [*] ya [=! hits the table imitating his teacher] .

%err:	palalos=pararos $PHO

%xepr:	$et3:GSA

*ADU:	y se enfada y cuanto <más grita más gritáis vosotros>

[=! deleted /s/ in final position of the word, dialectal] „ no?

*CHI:	0 [=! nods]. [ + trn]

%xepr:	$et3:GSA

*ADU:	porque no escucháis [=! deleted /s/, dialectal] [>].

*CHI:	<y: hoy:> [/] [<] y: hoy: [/] <hoy se ha> [/-] hoy [/-] y hoy se

[/] <se ha> [/] se ha cola(d)o@d en la claze [*] un paquete de

pañuelos [=! deleted /s/, dialectal] .

%err:	claze=clase $PHO

%xepr:	$i5:MMN:REP; $i5:MMN:REP; $i5:MMN:REF; $i5:MMN:REF; $i5:MMN:REP;

$i5:MMN:REP

*CHI:	y: [/] y después [=! aspirated /s/ in postnuclear position, deleted /s/ in final

position of the word, dialectal] el profe <lo ha> [/]

lo ha tira(d)o@d por la ventana [=! laughs].

%xepr:	$i5:MMN:REP; $i5:MMN:REP

*ADU:	muy bien .

*CHI:	<y tados> [*] [/-] [=! deleted /s/, dialectal] y todos

[=! deleted /s/, dialectal] azí [*] +"/.

%err:	tados=todos $PHO azí=así $PHO

%xepr:	$i5:MMN:REF;

*CHI:	+" buen viaje: [=! says goodbye with hand] !

%xepr:	$et3:GSA

*SPL:	pero que se ha colado dices ?

*CHI:	0 [=! nods] . [+ trn]

%xepr:	$et3:GSA

*MOT:	pero por qué l(o) ha tirado por la ventana ?

*CHI:	no sé [=! shrugs, laughs] .

%xepr:	$et3:GSA

*MOT:	un paquete de pañuelos ?

*ADU:	y sabes [=! deleted /s/, dialectal] cómo apareció el paquete de

pañuelos [=! deleted /s/, dialectal]?

*MOT:	porque <<estarían con e:l> [/] <estarían con el> [/-] estarían>

[=! aspirated /s/, dialectal] con <los pañuelos>

[=! deleted /s/, dialectal] y se habrá enfadao@d „ no ?

*CHI:	no [=! shakes head no] <estaba: [/] estaba>

[=! aspirated /s/, dialectal] en el suelo [=! shrugs].

%xepr:	$et3:GSA; $i5:MMN:REP; $et3:GSA

*CHI:	yo [/] yo <l(o) he visto> [?] coger [=! deleted /r/, dialectal] y:

lu [/-] [=! throw illustrator] allí (.) a:l <patio de infantil> [?] .

%xepr:	$i5:MMN:REP; $i5:MMN:REF; $et3:GSA

*MOT:	pues [=! deleted /s/, dialectal]

haberle [=! assimilation /r/ > /l/, dialectal] dicho +"/.

*MOT:	+" profe (.) no se tira !

*MOT:	y luego qué has [=! deleted /s/, dialectal] hecho „ hijo ?

*CHI:	<después [/] después> [=! aspirated /s/ in postnuclear position, deleted /s/ in final

position of the word, dialectal] otra vez me he ido a: la claze [*]

de apoyo <para la> [/] para la: [/-] e:h <para: la: xxx> [/-] para

la: cómo se llama ?

%err:	claze=clase $PHO

%xepr:	$i5:MMN:REP; $i5:MMN:REP; $i5:MMN:REF; $et3:DRA:FIL; $i5:MMN:REF;

$et3:DRA:ERQ

*CHI:	para: [/] <padá [*] [=! syllabified, regulator of rythm with hand]

hablar [=! deleted /r/, dialectal]> [/-] para <aprender a hablar>

[=! deleted /r/ in final position, dialectal] .

%err:	padá=para $PHO

%xepr:	$i5:MMN:REP; $et3:GSA; $i5:MMN:REF

*MOT:	<ah@i al [/-] la logopeda> [!] [>] has [=! deleted /s/, dialectal]

ido a la logopeda .

*CHI:	loope:da [*] [<].

%err:	loopeda=logopeda $PHO

*SPL:	muy bien .

*CHI:	y: la pro +/.

*MOT:	cómo se llama la: [/] la profe(sora) [/-] la logo(peda) +/?

*CHI:	Elena .

*MOT:	Elena.

*CHI:	y después [=! aspirated /s/ in postnuclear position, deleted /s/ in final

position of the word, dialectal] me he ido pada [*] claze [*] .

%err:	pada=para $PHO claze=clase $PHO

*CHI:	después [=! aspirated /s/ in postnuclear position, deleted /s/ in final

position of the word, dialectal] me [/] me he ido con una

zeño(rita) [*] y después [=! aspirated /s/ in postnuclear position, deleted /s/ in final

position of the word, dialectal]a caza [*] he comido +/.

%err:	zeño=seño $PHO caza=casa $PHO

%xepr:	$i5:MMN:REP;

*MOT:	pero no has [=! deleted /s/, dialectal] tenido recreo ?

*CHI:	he ido al lecleo [*] [=! nods] (.) después +/.

%err:	lecleo=recreo $PHO

%xepr:	$et3:GSA

*ADU:	y qué haces [=! deleted /s/, dialectal] en el recreo ?

*CHI:	jugar [=! deleted /r/, dialectal] al fulbo [*] .

%err:	fulbo=fútbol $PHO

*ADU:	al fútbol siempre ?

*CHI:	sí [=! nods] xxx sí (.) y también <<muchas veces> [/] muchas veces

nos> [=! deleted /s/, dialectal] toca <después del lecleo [*]> [/]

<despés [*] del lecleo [*]> [/] &desp [=! regulator with head] &cleo

toca: leligión [*] <y &po:> [/-] y: [/] y intan [*] <todo:s [/]

todos> [=! deleted /s/, dialectal] coriendo [*] para claze [*] y: ze

[*] pone un [/] un amigo mío +"/.

%err:	lecleo=recreo $PHO lecleo=recreo $PHO leligión=religión $PHO

intan=entran $PHO coriendo=corriendo $PHO claze=clase $PHO ze=se $PHO

%xepr:	$et3:GSA; $i5:MMN:REP; $i5:MMN:REP; $et3:DRA:DIS;

$et3:DRA:DIS; $i5:MMN:PER; $et3:GSA; $et3:DRA:DIS; $i5:MMN:REF;

$i5:MMN:REP; $i5:MMN:REP; $i5:MMN:REP

*CHI:	+" no: no corráis [=! deleted /s/, dialectal] que no: corráis

[=! deleted /s/, dialectal] <que lo> [/] <que lo> [/-] que nos

[=! deleted /s/, dialectal] toca con el profe: el Don+Juan podéis

[=! deleted /s/, dialectal] xxx allí !

%sit:	everybody laughs

%xepr:	$i5:MMN:REP; $i5:MMN:REF

@End
