## Supplemental file 1 for "Language impairment with a partial duplication of *DOCK8*"

**Inventory for Client and Agency Planning (ICAP)**

| **Adaptive Behavior Scales** | **RS** | **AE**  **Years- months** | **AE-CA**  **Years- months** | **ER**  **Years- months** | **DE** |
| --- | --- | --- | --- | --- | --- |
| Motor Skills | 486 | 5-10 | -4 - 11 | 4-7 to 8-0 | PER:1  TS: 52  RPI: 45/90 |
| Social and Communication Skills | 482 | 3-8 | -2 - 9 | 2-7 to 6-11 | PER:9  TS: 80  RPI: 73/90 |
| Personal Living Skills | 483 | 6-3 | +3 | 4-11 to 8-4 | PER:50  TS: 101  RPI: 91/90 |
| Community Living Skills | 489 | 8-1 | -2 - 5 | 6-8 to 10-0 | PER:10  TS: 81  RPI: 71/90 |
| Broad Independence (Total) | 485 | 6-6 | -2 - 9 | 4-7 a 8-10 | PER:9  TS: 80  RPI: 75/90 |
| **RS** (Raw score); **AE** (Age Equivalent); **CA** (Chronological Age); **ER** (Educational Rank); **DE** (Differential Score); **PER** (Percentile); **TS** (Typical Score); **RPI** (Relative Performance Index); | | | | | |

**Peabody Picture Vocabulary Test (PPVT-3)**

- Raw score: 69
- Scaled score:

‐ IC: 55

‐ Percentile: 1

‐ Eneatype: 1

**Test de Comprensión de Estructuras Gramaticales (CEG)**

| **Block** | **Type of sentence** | **Direct points (max. 4)** |
| --- | --- | --- |
| A | Predicative SVO (non-reversible)  E.g. El gato come un plátano ‘The cat eats a banana’ | 4/ |
| B | Attributive  E.g. El perro es negro ‘The dog is black’ | 4 |
| C | Negative predicative  El niño no come ‘The boy doesn’t eat’ | 4 |
| D | Pronominal predicative (with reflexive and non-reflexive pronouns)  E.g. La niña se lava las manos ‘The girl washes her (own) hands’  La mujer le pone los zapatos ‘The woman put him his shoes’ | 4 |
| E | Predicative SVO (reversible)  E.g. El ratón persigue al gato ‘The mouse chases the cat’ | 3 |
| F | Predicative SVO with a plural subject (reversible and non-reversible)  E.g. Los perros persiguen a la niña ‘The dogs chase the girl  Los niños ven la televisión ‘The boys watch TV’ | 4 |
| G | Coordinate disjunctive (with coordinated subject or object)  E.g. Ni el gato ni el perro son negros ‘Neither the cat nor the dog are black’  La niña no es ni rubia ni delgada ‘The girl is not either blond or thin’ | 4 |
| H | Predicative SVAdjunct (with *on*, *below*, *in front of*, or *behind*)  E.g. El perro está delante del gato ‘The dog is in front of the cat’ | 3 |
| I | Coordinate adversative (with coordinated subject or object and an intensifier)  E.g. No sólo el niño está jugando, sino también la niña ‘Not only the boy, but also the girl are playing’  La niña no sólo es rubia, sino también delgada ‘The girl is not just blond, but thin too’ | 4 |
| J | Relative (SO type)  E.g. El perro persigue al gato que es pequeño ‘The dog chases the small cat (lit. the cat that is small)’ | 4 |
| K | SVO with clefted subject  E.g. Es el gato el que muerde al perro ‘It is the cat that bites the dog’ | 4 |
| L | Comparative  E.g. El cuadrado es más grande que el círculo ‘The square is bigger tan the circle’ | 3 |
| M | OVS with focalized object  E.g. Al coche lo persigue la bicicleta ‘The car, the bike chases it’ | 1 |
| N | With a pronominalized object (contrasting only in gender)  E.g. Las niñas lo miran ‘The girls look at him’ | 4 |
| O | Relative (SS type)  E.g. El cuadrado que está dentro del círculo es azul ‘The square (that is) inside the circle is blue’ | 1 |
| P | Coordinate adversative (with coordinated subject or object and without an intensifier)  E.g. La niña es morena, pero el niño no ‘The girl is brunette, but the boy is not’  ‘The girl is thin, but not blond’ | 3 |
| Q | With a pronominalized object (contrasting both in number and gender)  E.g. El perro las persigue ‘The dog chases them (girls)’ | 3 |
| R | Passive OVS (reversible)  E.g. El gato es perseguido por el ratón ‘The cat is chased by the mouse’ | 2 |
| S | OVS with a clefted object  E.g. Es al ratón al que persigue el gato ‘It is the mouse that the cat chases’ | 0 |
| T | Relative (OS type)  E.g. El gato al que el perro persigue es pequeño ‘The cat that the dog chases is small’ | 3 |
|  | TOTAL | 62 (percentile 10^th^ ) |

**Registro Fonológico Inducido**

| ITEMS | Spontaneous speech | **Listen and repeat** |
| --- | --- | --- |
| 1. moto | + |  |
| 2. boca | + |  |
| 3. piña | + |  |
| 4. piano | + |  |
| 5. pala | + |  |
| 6. pie | + |  |
| 7. niño | + |  |
| 8. pan | + |  |
| 9. ojo | + |  |
| 10.llave | + |  |
| 11.luna | + |  |
| 12.campana | + |  |
| 13.indio | + |  |
| 14.toalla | + |  |
| 15.fuma | + |  |
| 16.dedo | + |  |
| 17.peine | + |  |
| 18.ducha | + |  |
| 19.gafas | + |  |
| 20.toro | + |  |
| 21.silla | + |  |
| 22.taza | + |  |
| 23.cuchara | + |  |
| 24.teléfono | + |  |
| 25.sol | + |  |
| 26.casa | + |  |
| 27.pez | + |  |
| 28.jaula | + |  |
| 29.zapato | + |  |
| 30.flan | + |  |
| 31.lápiz | + |  |
| 32.pistola | + |  |
| 33.mar | + |  |
| 34.caramelo | + |  |
| 35.plátano | + |  |
| 36.globo | + |  |
| 37.palmera | parmera |  |
| 38.clavo | + |  |
| 39.tortuga | totuga |  |
| 40.pueblo | + |  |
| 41.tambor | tambo |  |
| 42.escoba | + |  |
| 43.mariposa | + |  |
| 44.puerta | + |  |
| 45.bruja | bluja |  |
| 46.grifo | glifo |  |
| 47.jarra | + |  |
| 48.tren | + |  |
| 49.gorro | + |  |
| 50.rata | + |  |
| 51.cabra | + |  |
| 52.lavadora | + |  |
| 53.preso | preso |  |
| 54.semáforo | semaforo |  |
| 55.fresa | flesa |  |
| 56.árbol | arbol |  |
| 57.periódico | predriodico |  |
